# Normalization and De-noising of Single-cell Hi-C Data with BandNorm and 3DVI

**DOI:** 10.1101/2021.03.10.434870

**Authors:** Ye Zheng, Siqi Shen, Sündüz Keleş

**Author notes:** These authors have contributed equally (Ye Zheng, Siqi Shen). **Correspondence** Sündüz Keleş.

## Abstract

Single-cell high-throughput chromatin conformation capture methodologies (scHi-C) enable profiling long-range genomic interactions at the single-cell resolution; however, data from these technologies are prone to technical noise and bias that, when unaccounted for, hinder downstream analysis. Here we developed a fast band normalization approach, BandNorm, and a deep generative modeling framework, 3DVI, to explicitly account for scHi-C specific technical biases. We present robust performances of BandNorm and 3DVI compared to existing state-of-the-art methods. BandNorm is effective in separating cell types, identification of interaction features, and recovery of cell-cell relationship, whereas de-noising by 3DVI successfully enables 3D compartments and domains recovery, especially for rare cell types.

## Introduction

Maturation of chromosome conformation capture (3C) based technologies for profiling 3D genome organization^1–5^ and technological advancements in single-cell sequencing^6^ led to the development of single-cell Hi-C (scHi-C) assays^7–11^. Data from these assays enhance our ability to study the impact of spatial genome interactions on cell regulation at an unprecedented resolution. While some of the characteristics of scHi-C data, such as systematic genomic distance bias (referred to as *band bias*, Fig. 1A), are similar to its bulk version, scHi-C data harbors significantly distinct features. In general, data from single-cell technologies such as scRNA-seq and scATAC-seq are noisy and sparse, leading to underestimated biological signals within and across cells. However, these issues are compounded in 3C-based technologies because the natural analysis unit is locus-pairs depicting potentially interacting genomic loci; as a result, their sheer number exacerbates the sparsity. In contrast to bulk Hi-C data, which is available in small numbers of replicates owing to high sequencing cost, scHi-C is generated across thousands of cells simultaneously, therefore significantly increasing the resolution to capture biological variation. However, this resolution gain comes at the cost of increased technical noise and decreased sequencing depth per cell, further contributing to the extreme sparseness of scHi-C chromosomal contact matrices.

**Figure 1.**
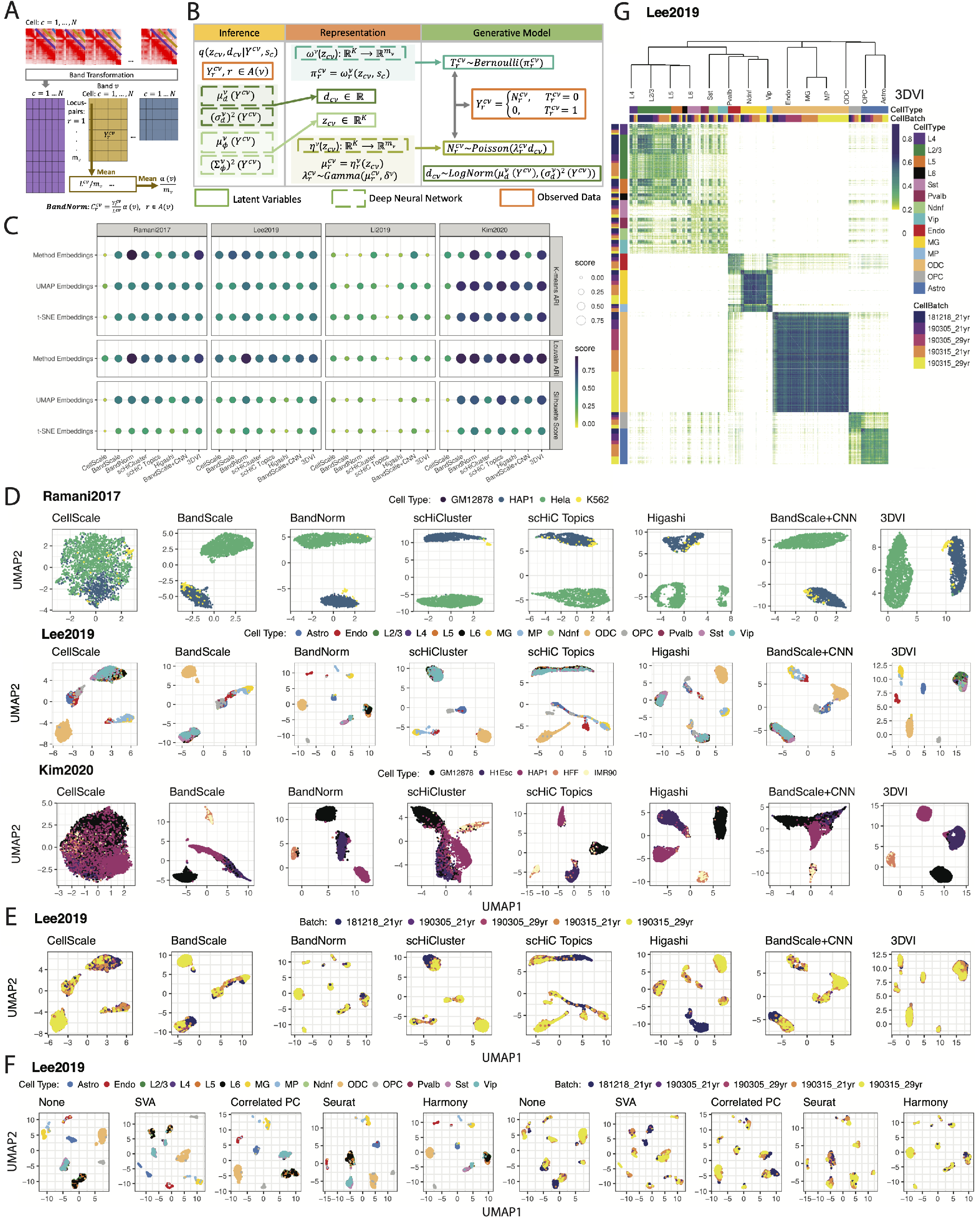
Benchmarking of scHi-C normalization and de-noising methods for celltype clustering. **A.** Band transformation separates scHi-C contact matrices into band-specific cell × locus-pair matrices before conducting **BandNorm** normalization on each band matrix per chromosome per cell. **B.** Deep generative model, **3DVI**, for a single band matrix *v* with locus-pair index set 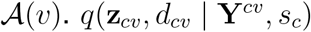 denotes the variational distribution of the latent factors, parameters of which are learnt locally for each band matrix. **C.** Evaluation of the eight scHi-C normalization and de-noising methods, namely CellScale, BandScale, BandNorm, scHiCluster, scHiC Topics, Higashi, BandScale+CNN, and 3DVI, for cell type separation across four benchmark datasets. The performances are evaluated by Adjusted Rand Index (ARI) after K-means clustering and Louvain graph clustering and by Silhouette coefficient on UMAP and t-SNE visualizations with the true cell labels. **D.** Application of the scHi-C data normalization and de-noising methods on `Ramani2017` with 4 cell types, `Lee2019` with 14 cells types, and `Kim2020` data sets with 5 cell types. The results are displayed using scatter plots of the two UMAP coordinates. The colors of the plotting symbols correspond to the cell types. **E.** Impact of batch effects on cell type separation using `Lee2019` data set with samples from two donors of ages 21 and 29 years old and in a total of 5 batches. The results are displayed using scatter plots of the two UMAP coordinates. The colors of the plotting symbols correspond to the batches. **F.** Results from the four batch effect removal methods, SVA^27^, removing the top correlated principal component, Seurat batch effect regression^28^, and Harmony^29^, together with the BandNorm normalization on the `Lee2019` data set. **G.** Pairwise cell similarity scores for the `Lee2019` data set. The scores are obtained by edge weights of the shared nearest neighbors graphs constructed on 3DVI latent embeddings of the data. The inferred relationship between the cell types is depicted by the dendrogram from the hierarchical clustering of the cells.

Initial approaches for unsupervised analysis of scHi-C data repurposed bulk data quantification methods of similarity between contact matrices, such as HiCRep^12^, and applied multidimensional scaling^13^. Vectorization of scHi-C contact matrices to form a cell by locus-pair matrix followed by dimension reduction approaches such as UMAP and t-SNE^11^, or topic modeling^14^ have been utilized successfully. Most recent approaches for normalization and denoising of scHi-C data rely on linear smoothing and random walk imputation^15^ of cell-specific contact matrices or hypergraph representation learning^16^. While these are highly innovative approaches, they lack a generative model that acknowledges the key properties of the scHi-C data. Deep generative modeling and, more specifically, variational autoencoders have seen a significant uptake in the analysis of single-cell transcriptomics^17,18^, epigenomics^19^, and proteomics^20^ due to their ability to provide efficient and scalable solutions to normalize, de-noise, and impute single-cell data. At the individual cell resolution, heterogeneity, driven by the stochastic nature of chromatin fiber and a multitude of nuclear processes, and unwanted variation due to sequencing depths and batch effects pose major analytical challenges for inferring single cell-level 3D genome organizations. Here we develop a deep generative model, named 3DVI, which systematically takes into account these structural properties and accounts for band bias, sequencing depth effect, zero inflation, sparsity impact, and batch effects of scHi-C data (Fig. 1B). In addition, we also describe a scaling normalization approach named BandNorm (Fig. 1A), and its variants as fast baseline alternatives (Methods).

## Results

We consider band transformation of scHi-C contact matrices, i.e., loci by loci symmetric matrices with entries representing the interaction frequency between the locus-pairs, as the foundation for our normalization and modeling approaches (Fig. 1A). The genomic distance effect, i.e., band effect, due to the random polymer looping behavior of DNA is one of the key features in both the bulk^1^ and single-cell Hi-C data (Supplementary Fig. 1). As expected, such a band effect leads to marked interaction frequency enrichment among loci close to the diagonal in the Hi-C contact matrix. Contact decay profiles that quantify interaction frequencies among locus-pairs as a function of their genomic distance can successfully separate cells based on their cell cycle stages^21^. To explicitly capture this effect, the upper triangular of the symmetric contact matrix for each cell is first stratified into diagonal bands, each representing a specific genomic distance between the interacting loci. Then, bands at the same genomic distance are combined into a band matrix across cells (Fig. 1A) for further BandNorm and 3DVI normalization (Methods). By dividing the interaction frequencies of each band within a cell with the cell-specific band mean, BandNorm first removes genomic distance bias within a cell and scales the sequencing depths between cells. Subsequently, BandNorm adds back a common band-dependent contact decay estimate by multiplying each band within a cell with the average band mean across cells (Fig. 1A and Methods). A similar strategy for imposing band-specific decay rates was adopted for normalization across bulk Hi-C samples^22^. In addition to this nonparametric normalization approach, we also devised 3DVI as a deep generative model built on the parametric count models of Poisson and Negative Binomial that have been successfully used in bulk measurements of chromatin conformation capture data^23,24^. Following the recent deep learning modeling approaches for single-cell transcription^17, 18^ and chromatin accessibility^19^, 3DVI builds a generative modeling framework on the band matrices for dimension reduction and de-noising of scHi-C data (Fig. 1B and Methods). It estimates and corrects the batch effect and de-noises interaction frequencies among locus-pairs that can then be leveraged for downstream analysis. BandNorm is implemented as an R package (https://github.com/keleslab/BandNorm) which also harbors all the curated public scHi-C data used in this paper. 3DVI is implemented as a python pipeline using the scvi-tools^17^ and is available at https://github.com/keleslab/3DVI.

We benchmarked BandNorm and 3DVI against two classes of methods, including baseline methods for library size and genomic distance effect normalization and more structured modeling approaches. In the former category, in addition to BandNorm, we evaluated CellScale and BandScale^11^ (Methods). CellScale uses a single scaling factor across all the locus-pairs within a cell, while BandScale employs a band-specific size factor, as opposed to a global one, within each cell to eliminate library size bias at each genomic distance^11^. After each of CellScale, BandScale, and BandNorm normalizations, single-cell contact matrices are vectorized into the cell by locus-pair matrices and used to generate low dimensional embeddings. Since this strategy overlooks the matrix structure of the data, we also utilized a convolutional neural network (CNN) approach, which has been previously leveraged for enhancing the resolution of the bulk-cell Hi-C matrix^25^, on contact matrices from BandScale to learn their low dimensional representations. In the second category, we considered three more state-of-the-art scHi-C methods: scHiCluster^15^, scHiC Topics^14^, and Higashi^16^.

We leveraged four scHi-C datasets with known cell-type labels and varying characteristics to evaluate the performances of the scHi-C methods (Methods). These four datasets are `Ramani2017`^9^ and `Kim2020`^14^ from various human cell lines, `Lee2019`^11^ from human brain prefrontal cortex cells, and `Li2019`^10^ from mESC cells. Of these, `Ramani2017`, `Lee2019` and `Kim2020` are generated from multiple sequencing libraries, hence, are especially suitable for investigating the impact of batch effects. All the data on chromosomes 1-22 and X are binned at 1Mb resolution to generate a set of loci, and raw data are filtered according to the specifications in the source publications to remove ultra sparse cells (Supplementary Table 1 and Supplementary Fig. 2).

We first assessed the cell-type separation performances of the scHi-C methods within the context of unsupervised clustering. We considered six evaluation settings (Fig. 1C and Methods): K-means clustering of the latent embeddings of each method, and with or without low-dimensional projections with t-SNE and UMAP, and Louvain graph clustering^26^. We used Adjusted Rand Index (ARI) to compare the resulting clusters with the known cell labels and the average Silhouette score to quantify the separation between the clusters. Cell clustering performances vary dramatically due to the numbers of cells, data quality as measured by sequencing depth, sparsity, and batch effects, as well as how distinguishable the cell types are. For the `Ramani2017` and `Lee2019` studies, BandNorm and 3DVI perform as well as or outperform the rest based on all the six evaluation settings (Fig. 1C). Almost all the methods perform poorly on the `Li2019` dataset (ARI ∈ [0.005, 0.47] and Silhouette score ∈ [−0.56,0.2]), which has the smallest number of cells (Supplementary Table. 1 and Supplementary Fig. 2). All the methods except CellScale achieve their best performances on the Kim2019 dataset (ARI ∈ [0.35, 0.91] and Silhouette score ∈ [0.3, 0.73]), with leading performances by BandNorm, Higashi, scHiC Topics, and 3DVI (Fig. 1C). Overall summary of these evaluations yields Band-Norm and 3DVI as robustly best performing, followed by scHiCluster and Higashi, with median rank scores of 2, 2.5, 4, and 4.5, respectively (Supplementary Fig. 3). In addition to these quantitative measures, we also graphically assessed whether the resulting low-dimensional representations achieve clear cell type separation (Fig. 1D), and carry left-over effects by technical variation due to batch (Fig. 1E), library size or sparsity (Supplementary Fig. 4). We present `Ramani2017, Lee2019` and `Kim2020` with UMAP visualizations for illustration (Fig. 1D) and provide the t-SNE embeddings (Supplementary Figs. 5-7) as well as analysis for `Li2019` (Supplementary Figs. 8 and 9) in Supplementary materials. More specifically, across all four datasets, CellScale, a commonly used global library size scaling strategy for high-throughput sequencing data, produces the least favorable cell separation. Particularly, for `Ramani2017` and `Kim2020` (Fig. 1D) where most of the other methods can explicitly separate major cell types, CellScale exhibits no separation power. For `Ramani2017` (Fig. 1D), BandNorm and scHiCluster stand out in successfully separating the small numbers of K562 cells from the rest. Higashi, on the other hand, exhibits batch effects resulting in the further division of the Hela cells into sub-clusters (Fig. 1D and Supplementary Figs. 5 and 10). The original analysis of `Lee2019` reported that scHi-C contact matrix data alone could barely separate the excitatory neuronal subtypes, namely L2/3, L4, L5, and L6, from the inhibitory cells Ndnf, Pvalb, Sst, and Vip. While this is the case for most methods, BandNorm and 3DVI show a notable exception. Specifically, BandNorm unambiguously isolates the excitatory and inhibitory cells into two clusters. Consistent with this, 3DVI also achieves a clear boundary between the excitatory and inhibitory cells (Fig. 1D). Direct comparison of the results from BandScale to BandNorm highlights the marked improvement due to adding back the band-specific contact decay estimates by BandNorm (Figs. 1A, D, and Methods). Allowing band-specific contact decays can be interpreted as adding more weights to short-range bands and plays a significant role in separating the cell types. Higashi analysis of the `Lee2019` data, however, also shows nontrivial batch and sequencing depths effect, leading to erroneous separation of the mixture of the neuron subtypes (L2/3, L4, L5, and L6) and inhibitory cells (Ndnf, Pvalb, Sst and Vip) into two mixture clusters (Figs. 1D, E, Supplementary Figs. 4 and 6). scHiCluster requires stringent cell filtering (Table 1 and Methods) that may remove 88.4% of cells in the dataset^14^. Without such stringent filtering, scHiCluster loses its cell type separation power unexpectedly. For example, in the `Kim2020` study where BandNorm and other more structured models achieve distinct separation with high ARI and Silhouette scores (Figs. 1C, D and Supplementary Fig. 7), scHiCluster, performs poorly in distinguishing HAP1 and H1Esc. This is consistent with the observations of others^14,16^ on scHiCluster. While scHiC Topics performs impressively well on `Kim2020`, it does not exhibit similarly high performance on the other datasets. This is in spite of the fact that we used the suggested strategy^14^ of setting the number of topics based on the true cell labels in the Silhouette score calculations. The BandScale+CNN strategy explicitly acknowledges the matrix structure of the data and can potentially learn the graph structure of each cell matrix. However, it yields limited power, possibly owing to the sparsity and low resolution of 1Mb. 3DVI is generally performing as well as BandNorm in terms of separating cell types, and its overall high performance is stable across datasets and robust to batch effects (Fig. 1E and Supplementary Fig. 10).

**Table 1:** Overview of single-cell Hi-C data analysis methods.

| Method | Resolution | Cell Filtering | Locus-pairs Filtering | Sequencing Depth Normalization* | Batch Effect Removal | Time (Lee2019) |
| --- | --- | --- | --- | --- | --- | --- |
| CellScale | 1Mb | NULL | NULL | Each locus-pair is normalized by dividing with the total IFs of each contact matrix and multiplying by 10,000. | NULL | <15min(23cores) |
| BandScale<sup>[11]</sup> | 100kb | NULL | Keep $\leq 2$Mb | Each locus-pair is normalized by dividing with the average IFs of locus-pairs at the same distance. | NULL | <15min(23cores) |
| BandNorm | 1Mb | NULL | NULL | Each locus-pair is normalized by dividing with the total IFs at the same genomic distance (i.e., band) per cell and multiplying by the average total interaction frequency across cells at the same distance. | Top batch correlated PCA components other than the 1st and 2nd are removed. | <15min(23cores) |
| scHiCluster<sup>[15]</sup> | 1Mb<br>200kb | Keep cell with off-diagonal contacts $> 5000$. | Keep cells with $\geq x$ non-zero locus-pairs, $x$ is chromosome size (Mb). | After neighborhood smoothing and random walk imputation, top 20% interacting locus-pairs are set to 1 and rest to 0. | NULL | 30min(23cores) |
| scHiC Topics<sup>[14]</sup> | 500kb | NULL | Keep $\leq 10$Mb | Cell by locus-pair matrix is binarized; (Optional) Bottom 50% cell based on total IFs are removed and the rest is downsampled to remove the sequencing depth effect. | NULL | 36h(1core) |
| Higashi<sup>[16]</sup> | 1Mb | Keep cells with $\geq 2,000$ interactions that have genomic span greater than 500Kb. | NULL | Contact maps can be smoothed via linear convolution to reach similar sparsity; Quantile normalization can be applied to the matrix. | NULL | 49h(10coresCPU)<br>16h(23coresGPU) |
| BandScale+CNN | 1Mb | NULL | Keep cells with $\geq x/6$ non-zero locus-pairs, $x$ is chromosome size (Mb). | Each locus-pair is normalized by dividing with the average IFs of locus-pairs at the same distance. | NULL | 5h(23cores) |
| 3DVI | 1Mb | NULL | (Optional) Diagonals of contact matrix with high contact count variation are selected. | A variational autoencoder with Gamma-Poisson mixture at each band with a separate size factor is fitted. | The batch factor is incorporated into the autoencoder at each distance. | 4h(23coresGPU) |

\*: IF refers to interaction frequency.

Batch effects emerge as a key modulator of the cell type separation performances of different methods. Specifically, visualization based on low-dimensional embeddings of the methods indicates Higashi as affected most, followed by scHiC Topics, scHiCluster, and BandScale+CNN (Fig. 1E and Supplementary Fig. 10). 3DVI explicitly models the batch factor into the deep generative framework hence does not show severe batch biases. BandNorm normalization step only considers the distance effect and library size without any adjustment for the batch factor. Therefore we observed unexpected cell separation within excitatory neuronal subtypes of `Lee2019` (L2/3, L4, L5, L6, Fig. 1F-None panel). In order to address this, we considered several methods that yielded promising results in removing batch effects for other types of high-throughput sequencing data, including SVA^27^, removing the principal component exhibiting the highest correlation with the batch variable, Seurat batch effect regression^28^, and Harmony^29^. Of these, SVA^27^ and removing the principal component exhibiting the highest correlation with the batch variable do not remove the batch bias, while SVA even makes the performance worse. Seurat batch effect regression^28^ alleviates the separation within the excitatory neuronal subtypes cluster but introduces additional batch biases (depicted in orange and yellow of Fig. 1F) for other clusters. Harmony^29^ stands out in successfully addressing the batch effects and enhances the sub-cell type separation. Additionally, for `Ramani2017` data, with five libraries where ml1 and ml2, and pl1 and pl2 are sequencing experiments with same library preparations, all four batch removal methods work successfully in mixing pl1 library with its pair pl2 (Supplementary Fig. 11A). For `Kim2020`, however, the libraries are confounded with the cell type; therefore, it is not suitable to apply any batch removal strategy, and doing so worsens the cell type separation (Supplementary Fig. 11B). In addition to confounding cell-type separation, batch effects may also impact inference of cell type relationships from the pairwise similarity measurements of the cells (Fig. 1G). We compared the robustness of the methods regarding this effect by considering a cell-to-cell similarity metric based on the edge weights of shared nearest neighbor graphs^30^. As revealed by the hierarchical clustering of the cells based on this metric (Fig. 1G and Supplementary Fig. 12), CellScale, BandNorm, scHiCluster, scHiC Topics, Higashi, and 3DVI latent embeddings successfully separate the excitatory neuron subtypes (L2/3, L4, L5, and L6), CGE-derived inhibitory subtypes (Ndnf and Vip), medial ganglionic eminence-derived inhibitory subtypes (Pvalb and Sst), oligodendrocyte related cell types (Astro, OPC, ODC) and non-neuronal cell types (MG, MP, Endo). BandNorm after Harmony batch removal yields the most invariant performance to the batch effects on the similarity matrix (Supplementary Fig. 12).

Another notable observation from this benchmarking study is that UMAP and t-SNE visualizations of low-dimensional embeddings from BandScale, scHiCluster, scHiC Topics, Higashi, BandScale + CNN, and 3DVI tend to display systematic effects of sequencing depth and sparsity despite their implicit or explicit efforts to account for these factors (Supplementary Fig. 4). However, these lingering effects do not impact the overall cell type separation. Instead, they tend to impact the local organization of the cells within their respective clusters. We considered utilizing PCA on the low-dimensional embeddings (t-SNE or UMAP) to identify and remove the top principal component(s) highly correlated with sequencing depth and sparsity. However, their removal did not result in any discernible improvement in the cell type separation and, on the contrary, led to worse evaluation metrics for some datasets. This eludes to the possibility that the sequencing depth and sparsity effects left in the model embeddings are confounded with other latent biological or technical variation hence cannot be completely removed from the data.

Next, we sought to evaluate the normalization and de-noising by BandNorm and 3DVI along with other methods for their impact on downstream scHi-C data analysis, leveraging the `Kim2020` dataset as an illustration (Fig. 2). We compared aggregated scHi-C contact matrices of individual cell types after normalization (BandNorm) or de-noising (3DVI, Higashi, and scHiCluster) with their existing bulk Hi-C versions as the gold standard in terms of detection of A/B compartments and topologically associating domains (TADs), contact matrix similarity, detection of significantly interacting and differentially interacting locus-pairs. The results presented are for scHi-C aggregation based on the true cell type labels; however, the overall benchmarking conclusions remain the same for the aggregation using unsupervised clustering labels (Methods). When the number of cells per cell type is large such as in GM12878 (Supplementary Fig. 2), BandNorm normalized and aggregated scHi-C data exhibits good visual agreement with the bulk version in terms of the broad features of the contact matrix, such as the TADs and A/B compartments (Fig. 2A). However, for rare cell types, i.e., IMR90 in the `Kim2020` dataset, data from BandNorm reflects extreme sparsity and does not yield good concordance with the bulk version (Fig. 2B). In contrast, contact matrices de-noised with 3DVI appear more in agreement with their bulk version regardless of the numbers of cells (Figs. 2A, B). Higashi borrows information from the neighboring cells in a hypergraph embedding for imputation. This may lead to over-imputation, as in the case of IMR90 cells (Fig. 2B). scHiCluster de-noises the scHi-C contact matrix through neighborhood smoothing and random walk; hence the matrices look smooth and blurry compared to their bulk versions. Systematic quantification of these observations indicates their generality and consistency across the cell types. At the domain structure level, despite the sparsity in the aggregated scHi-C matrices, BandNorm has the most concordant insulation score (purple lines in the “Insulation Score” panel of Figs. 2A, B, Supplementary Fig. 13) to that of bulk data (grey lines in the “Insulation Score” panel of Figs. 2A, B, Supplementary Fig. 13). This results in the highest recovery rate for TAD boundaries across all the chromosomes and all the five cell types (Fig. 2C, e.g., the median recovery rates for GM12878 are 85.71%, 66.67%, 63.64% and 57.14% for BandNorm, 3DVI, Higashi and scHiCluster, respectively). Furthermore, HiCRep^12^ similarity analysis confirms that Band-Norm normalization has the overall highest reproducibility score with the bulk Hi-C data, followed by 3DVI and Higashi (Fig. 2D). Surprisingly, pairwise cell line reproducibility analysis reveals a strong correlation between HFF and IMR90 contact matrices after imputation by Higashi (Fig. 2E). Further investigation attributes this to Higashi’s imputation strategy whereby it borrows information from neighboring cells and relies strongly on the accurate separation of the cells based on their cell types to form informative neighborhoods. As revealed by the UMAP and t-SNE visualization of cell type clustering (Fig. 1D and Supplementary Fig. 5), none of the methods can isolate IMR90 cells, and most IMR90 cells are mixed with the HFF cell cluster. As a result, neighbors of IMR90 cells inevitably consist of cells from a variety of cell types and are especially enriched in HFF cells. This explains the high similarity between IMR90 and the other four aggregated scHi-C cell types after Higashi imputation, particularly for the HFF cell line (Fig. 2E). This observation again underlines the concerns of over-imputation for Higashi as observed visually on the contact matrices (Figs. 2A, B, Supplementary Fig. 13).

**Figure 2.**
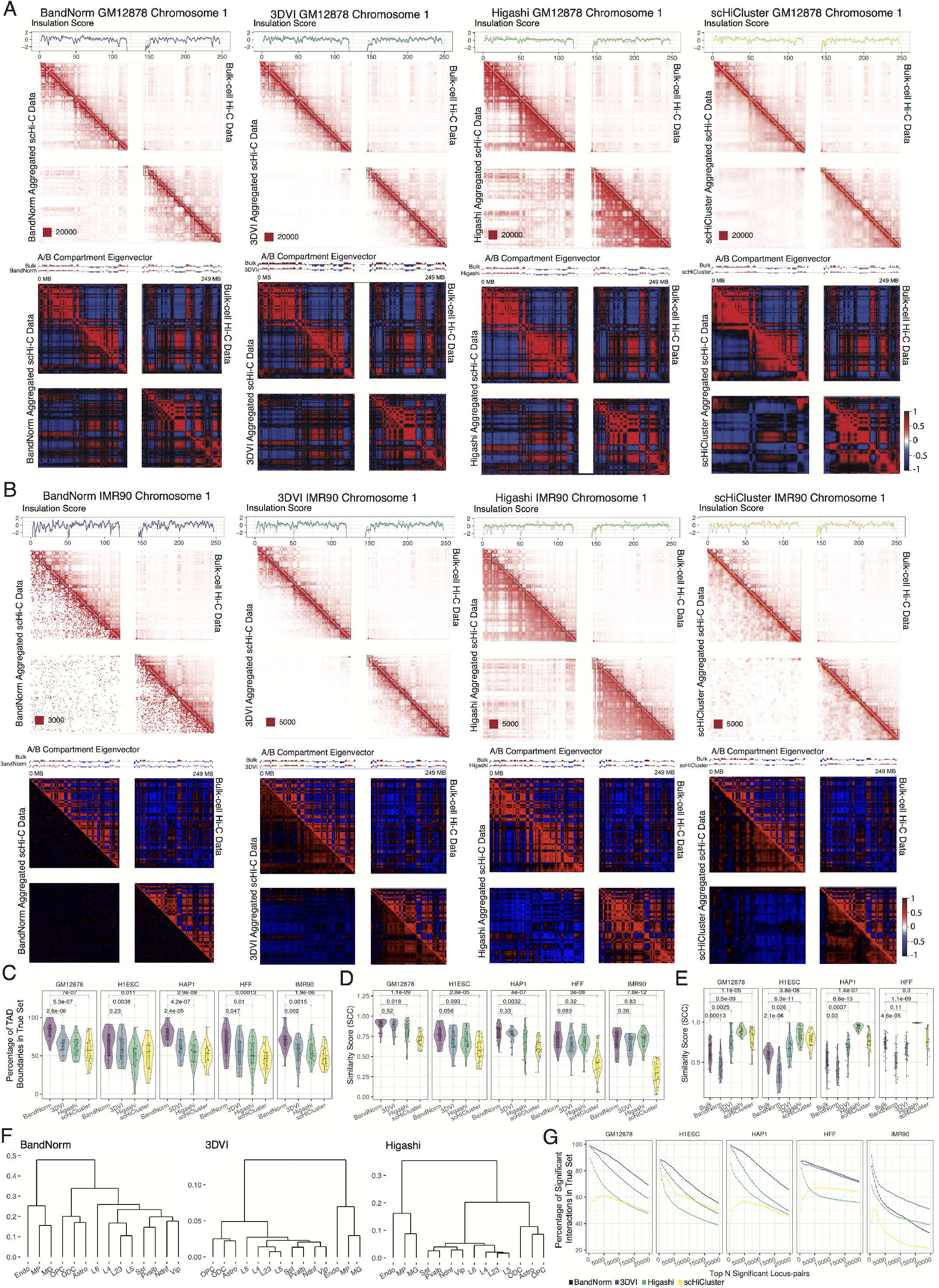
Evaluation of scHi-C normalization and de-noising methods for their impact on the downstream Hi-C data analysis. **A.** Comparison of detected topologically associating domains (TADs) and A/B compartments between bulk GM12878 Hi-C data on chromosome 1 (upper right triangles) and the aggregated single-cell Hi-C data (lower left triangles) after normalization or de-noising using `Kim2020` data set. True cell labels for GM12878 were utilized to aggregate processed scHi-C data. First row: The insulation scores^36^ that trace the TAD boundaries are depicted on the contact matrices with grey lines corresponding to bulk Hi-C data and purple for BandNorm, blue for 3DVI, green for Higashi, and yellow for scHiCluster. The numbers after the red squares at the left bottom of each contact matrix represent the minimum interaction frequency for the reddest locus-pair. Second row: A/B compartments are detected using the eigenvector of correlation map of bulk (upper right triangles) or aggregated (lower left triangles) Hi-C matrices, values of which are displayed above each correlation matrix. **B.** Comparison of detected TADs and A/B compartments between bulk IMR90 Hi-C data and the aggregated single-cell Hi-C data after normalization or de-noising using `Kim2020` data set with the known IMR90 cell type label. **C.** Percentage of TAD boundaries that are within 1Mb distance of the corresponding bulk cell type Hi-C data TAD boundaries. The numbers above each pairwise comparison with BandNorm are the *P* values based on the two-sided t-test. **D.** HiCRep^12^ similarity scores between aggregated scHi-C matrices and the corresponding bulk Hi-C matrices. The numbers above each pairwise comparison with BandNorm are the P values based on the twosided t-test. **E.** HiCRep^12^ similarity scores between aggregated IMR90 matrices and aggregated GM12878, H1Esc, HAP1, and HFF from `Kim2020` data set. The numbers above each pairwise comparison with bulk are the P values based on the two-sided t-test. Sample sizes for **C-E** are n = 23 for each violin corresponding to chromosome 1-22 and chromosome X. **F.** Hierarchical clustering of the aggregated scHi-C matrices of different cell types from BandNorm, 3DVI, and Higashi based on their HiCRep similarity scores. **G.** Percentage of top N (N = 5,000, 10,000,…) significant interacting locus-pairs that are in the gold standard set for each method. The gold standard set is defined as the top 50,000 significant locus-pairs detected by Fit-Hi-C^31^ from the cell type specific bulk Hi-C data.

A use case of scHi-C data is the inference of similarity between different cell types based on their 3D chromatin interactions. We first evaluated the hierarchical relationship of the aggregated scHi-C normalized or de-noised matrices across 14 cell types of `Lee2019` dataset constructed based on the HiCRep similarity scores (Fig. 2F and Supplementary Fig. 15A). Alternatively, we also considered similarity at the individual cell level and quantified cell-to-cell similarity based on the edge weights of the shared nearest neighbor graphs on the matrices de-noised by individual methods (Supplementary Fig. 14). For all the methods, both the aggregated and single cell-level similarity measurements demonstrated a clear separation of neural sub-cell type clusters as suggested in the literature^11^. Notably, aggregated scHi-C data from BandNorm, 3DVI, and Higashi yielded better segregation of the endothelial cells (Endo) compared to scHiCluster.

We next investigated the performances of the de-noising methods at the locus-pair level in terms of detection of significantly interacting locus-pairs within a cell type and differentially interacting locus-pairs across cell types. We first set a gold standard using the cell line specific bulk Hi-C data. Specifically, we identified the top 50,000 significant interacting locus-pairs from bulk Hi-C with Fit-Hi-C^31^ as the “true” significant interaction list. Then, we quantified the percentage of this list that can be recovered by the top interacting locus-pairs from the aggregated scHi-C matrices of each method. BandNorm outperforms all the other methods by achieving the highest accuracy rate (e.g., for the top 5000 interacting locus-pairs of GM12878, the accuracy rates vary as 93.34%, 83.32%, 68.36% and 61.10% for BandNorm, 3DVI, Higashi, and ScHiCluster, respectively), across all cell lines except for the IMR90 scHi-C data which has the smallest number of cells (Fig. 2G). 3DVI maintains a highly significant interaction detection accuracy rate for the IMR90 cells owing to its successful imputation strategy (Fig. 2G). In order to evaluate performances for detection of differential TADs or locus-pairs, we considered TADcompare^32^ for differential TAD boundaries detection, diffHic^33^ for differential interacting locus-pairs detection, and CHESS^34^ for differential interacting regions detection (Supplementary Fig. 15). Overall, aggregated scHi-C matrices from all four methods resulted in similar differential interaction detection.

Finally, we quantified the run time of each method on the `Lee2019` dataset which has large numbers of cells with high sequencing depths. Higashi experiments using GPU version and 3DVI were carried out on a machine with 18 cores 2 sockets Intel(R) Xeon(R) Gold 6254 CPU addressing 562GB RAM and one NVIDIA GeForce RTX 2080 Ti GPU addressing 11GB RAM. The rest of the methods, including Higashi experiments only using CPU, were tested on a machine with 14 cores 2 sockets Intel(R) Xeon(R) CPU (E5-2680 v4) addressing 256GB RAM. The run times of the methods vary dramatically from less than 15 minutes to several hours or even days using multiple cores for parallel running with one core per chromosome (Table 1).

The profiling of single-cell 3D genome organization is poised to generate new types of investigations at unprecedented and unique levels of resolution. A key analytical task of these investigations is de-noising and normalization of scHi-C data to infer clusters of cells representing cell types, stages, and conditions. We presented BandNorm and 3DVI at the two opposite ends of the structured modeling of scHi-C data. Our evaluations on datasets with known cell types and varying data characteristics indicate that BandNorm, which corrects for genomic distance bias and library size, performs as well and even better than elaborate modeling approaches on these datasets. When coupled with Harmony, BandNorm is also robust to batch effects in cell type separation. In comparison, 3DVI can account for genomic distance, library size, sparsity, and batch effects and impute sparse contacts to provide advantages for downstream analysis compared to Bandnorm. We note that the current version of 3DVI does not consider spatial dependence among adjacent locus-pairs, i.e., (*i,j*) with (*i* — 1, *j* — 1) and (*i* + 1, *j* + 1) in the band matrices. While high-resolution contact maps, e.g., locus size ≤ 10Kb from bulk Hi-C data exhibit local spatial dependency between interacting loci^35^, the spatial independence assumption is well justified for low resolution (200Kb-1Mb) scHi-C data (Supplementary Fig. 16).

## Acknowledgments

This work was supported by NIH grants HG003747 and HG011371 to S.K. We thank Dr. Peigen Zhou, Fan Chen and Shan Lu from University of Wisconsin - Madison for insightful discussions.

## Author Contributions

S.K. conceived the project. Y.Z., S.S. and S.K. designed the research and developed the methods. All the authors contributed to the preparation of the manuscript.

## Methods

### Band transformation of scHi-C data

Band transformation of scHi-C contact matrices forms the basis for our normalization and modeling approaches. To explicitly capture the genomic distance effect, the upper triangular of the symmetric contact matrix for each cell is first stratified into diagonal bands, each representing the genomic distance between the interacting loci. Then, bands from the same genomic distance are combined into a band matrix across cells (Fig. 1A). Specifically, we denote the set of bands by 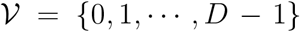, where *D* denotes the number of loci in the contact matrix (i.e., # of rows) and *v* = 0 represents the diagonal band, 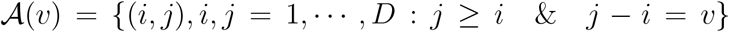 indices of the locus-pairs in the band *v*, and *m_v_* the total number of locus-pairs in the band v. For the modeling and downstream analysis, we only consider the off-diagonal interactions, hence *v* ≥ 1.

### BandNorm: A fast baseline band normalization approach for scHi-C data

Bandnorm provides fast normalization of scHi-C data by first removing genomic distance bias within a cell and sequencing depth normalizing between cells followed by adding back a common band-dependent contact decay estimate for the contact matrices across cells. More specifically, the BandNorm normalized interaction frequencies are obtained by

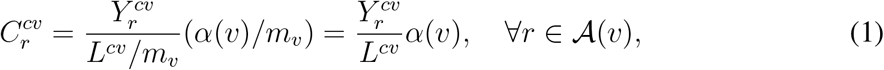

where *L^cv^* denotes the total interaction frequency of cell *c* at the *v*-th band, 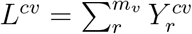, and *α*(*v*) is the average interaction frequency of band v across cells and is defined as 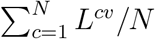.

### 3DVI: A deep generative model for scHi-C data

3DVI models the interaction frequencies of locus-pairs in each cell as a sample from a zero-inflated negative binomial distribution while accounting for library size and batch effect for each band matrix. Let 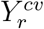 denote the interaction frequencies, i.e., quantification of how strongly two loci interact, between genomic loci *i* and *j* where 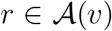 in cell *c* = 1,…, *N* at band *v* and Y^*cv*^ vector of interaction frequencies of locus-pairs for cell *c* at band *v*. The two key components of this model are a non-linear latent factor model to obtain low-dimensional representations z_*cv*_ of cell c across band matrices Y^*cv*^ and a hierarchical generative model for Pr(Y^*cv*^ | z_*cv*_). Specifically, we model observed interaction frequencies 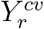 for each locus-pair in Y^*cv*^ with a latent space model. Low dimensional latent variable z_*cv*_ enables nonlinear dimension reduction for characterizing differences among cells in band *v*. Next, we present the generative process where each interaction frequency 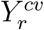 is drawn independently conditional on z_*cv*_ through the following process, where we assume that z_*cv*_ ~ Normal(0, *I_K_*). In order to account for the sparsity of the scHi-C data, we define a zero inflation variable 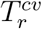 and set 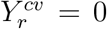 if 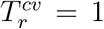 and 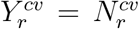 otherwise. Here, 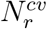 denotes the observed interaction frequency in the absence of a dropout, and 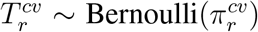, where 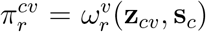. s_*c*_ is the batch information for cell *c* and *ω*(.) is a neural network that encodes whether a particular locus-pair has dropped out due to the technical artifacts^23^ and maps the latent space z_*cv*_ back to the full dimension of all locus-pairs in band *v*. Additionally, 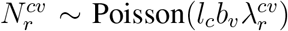 where 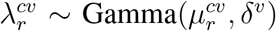, *l_c_* denotes the latent library size factor for cell *c*, and the band size factor *b_v_* modulates the impact of the size factor on the true interaction frequencies for band *v*. The band size factor, *b_v_*, is motivated by the genomic distance effect in which interaction frequencies between locus-pairs vary systematically by the distance between the loci. The band effect has been observed to vary markedly between two bulk Hi-C experiments^24^. We inquired whether this effect varied depending on the observed library size of the individual cells by leveraging one of our case study data sets `(Lee2019)` and formally tested for an interaction between library size and band. Specifically, we considered a mixed linear model for cell-specific mean band interaction frequencies as a function of cell type, observed library size, and band indices as

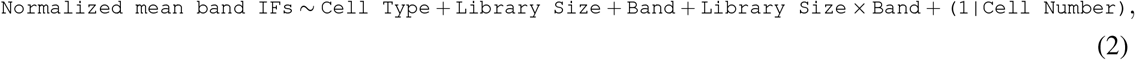

where interaction frequencies are normalized to per million within each cell and log transformed, and the model term `(1|Cell Number)` denotes the random effects of the cells to accommodate potential correlations between measurements from the same cell. This analysis revealed a significant interaction between library size and band (*P* value << 10e — 6), and the model with the interaction term fitted better than the smaller submodels based on the Bayesian Information Criterion (BIC^37^). This observation enables a more flexible parametrization of scaling factor that merges library (*l_c_*) and band size (*b_v_*) factors into cell type specific band size factors as *d_cv_*. Moreover, we let 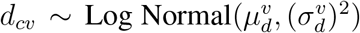 parametrize the prior for this scaling factor in the generative model.

Next, we modeled the mean parameter 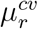 as a nonlinear function of z_*cv*_ as 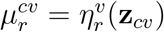, where η(.) is a neural network that maps the latent space to the full dimension of the locus-pairs. We leveraged existing variational inference tools, scvi-tools (single-cell variational inference tools^17^), to fit this model for each chromosome separately. The estimated latent components z_*cv*_ of each cell were concatenated across bands and chromosomes for final low dimensional projection by UMAP or t-SNE and downstream analysis.

### scHi-C data analysis methods compared in the benchmarking experiments

In our evaluations of unsupervised clustering of the cells based on scHi-C data, we considered two classes of methods, including baseline methods for library size and genomic distance effect normalization and more structured modeling approaches. In the former category, in addition to BandNorm, we evaluated CellScale, which uses a single scaling factor across all the locus-pairs within a cell and scales library sizes of each cell to a common size (10,000 in our applications). We also considered the first normalization step of BandNorm separately as BandScale, where interaction frequencies of each band within a cell are divided by the cell’s band mean. BandScale uses band-specific size factors rather than a global size factor within a cell and has been used previously to eliminate library size bias at each genomic distance^11^. After each of CellScale, BandScale, and BandNorm normalizations, single-cell contact matrices are vectorized into the cell by locus-pair matrices and used to generate low dimensional embeddings. To incorporate the matrix structure of the data, we utilized a convolutional neural network (CNN) approach, which has been previously leveraged for enhancing the resolution of the bulk-cell Hi-C matrix^25^, on contact matrices from BandScale to learn the lower-dimensional representation of the contact matrices.

Among the more structured modeling approaches, in addition to 3DVI, we also considered the state-of-the-art scHi-C data processing methods scHiCluster, scHiC Topics, and Higashi. scHiCluster starts with neighborhood smoothing and random walk imputation to reduce the contact matrix sparsity. Then, it binarizes the imputed matrix by setting the top 20% interacting locus-pairs to 1 and the rest to 0 in order to remove the library size bias across cells. For dimension reduction, the contact matrix of each chromosome is vectorized into a cell by locus-pair matrix, and the top 20 PCA components are concatenated across chromosomes for a second PCA run. The final top principal components are used for clustering and visualization. Notably, scHiCluster requires the most stringent cell filtering where only cells with total off-diagonal interaction frequencies > 5,000 are kept. It further enforces less sparsity by discarding cells with less than *x* non-zero locus-pairs, where *x* is the number of *x* Mb loci on each chromosome, separately for the contact matrices of each chromosome. scHiC Topics focuses more on the short to mid-range interactions by only considering the intra-chromosomal locus-pairs within 10Mb genomics distance. This aims to balance the data sparsity and reduce model complexity. Contact matrices of the cells are first vectorized to construct the cells by locus-pairs matrix, which is further binarized and input into a topic modeling framework. Cell type clustering is implemented on the cell by topics matrix where “topics” are proxies for cell types. Higashi trains a hypergraph neural network and enables neighboring cells in the hypergraph to share information for capturing interaction patterns. The resulting embeddings are then used for learning cell types. Table 1 summarizes these eight methods in terms of their pre-processing and treatment of various sources of biases.

### Benchmark datasets

We considered four existing studies with varying data characteristics and known cell types to benchmark the scHi-C low-dimensional embedding approaches. These four datasets are scHi-C measurements from human cell lines (`Ramani2017`^9^ and `Kim2020`^14^), human brain prefrontal cortex cells (`Lee2019`^11^), and mESC cells (`Li2019`^10^).

`Ramani2017` has four human cell lines, HeLa S3, HAP1, K562, and GM12878. These cell lines are distributed over five sequencing libraries labeled as ml1, ml2, ml3, pl1, pl2, where pairs ml1 and ml2, and pl1 and pl2 are sequencing experiments with the same library preparations, respectively, and hence present different batches. We downloaded the `Ramani2017` data from GEO^38^ with data accession GSE84920, and followed the instructions of literature^9^ to filter out low count reads. Data was transformed into the sparse matrix format at 1Mb by scHiCTools^39^.

`Lee2019` generated scHi-C data from 14 human brain prefrontal cortex cell types, Astro, Endo, L2/3, L4, L5, L6, MG, MP, Ndnf, ODC, OPC, Pvalb, Sst, Vip, originating from two donors with ages of 21 and 29 years and in a total of five sequencing libraries. Data were downloaded from https://salkinstitute.app.box.com/s/fp63a4j36m5k255dhje3zcj5kfuzkyj1/folder/82405563291. This dataset has the largest numbers of sequencing reads and the highest average interaction frequency per cell. Furthermore, since all the cells are prefrontal cortex cells, they are expected to exhibit less heterogeneity compared to other datasets.

`Li2019` datasets harbor mESC cells cultured in serum and leukemia inhibitory factor (LIF) condition (serum mESCs: serum 1 and serum 2) and mESCs cultured in LIF with GSK3 and MEK inhibitors (2i) condition. This data is valuable in benchmarking the performances of the methods when the number of cells is small. We downloaded the `Li2019` data from http://enhancer.sdsc.edu/ligq/share/mESC.MH/scMH/juice_files/ and converted into sparse contact matrices by Juicer^40^.

`Kim2020` dataset contains scHi-C data from five human cell lines, GM12878, H1Esc, HAP1, HFF, and IMR90, with nine sequencing libraries. While this data has the largest number of cells, the average off-diagonal interaction frequency per cell is the smallest. Notably, the numbers of cells vary dramatically across cell types, with GM12878, H1Esc, and HAP1 having more than 2k cells and IMR90 with less than 100 cells (Supplementary Fig. 2). We downloaded the data from https://noble.gs.washington.edu/proj/schic-topic-model/.

All the scHi-C data on chromosomes 1-22 and chromosome X were binned at 1Mb resolution to generate a set of loci, and extremely sparse cells were removed if the number of non-zero locus-pairs was less than *x*/6 for the contact matrix of each chromosome where *x* is the chromosome size in Mb (Supplementary Table 1). We discarded the scHiCluster cell filtering requirement (Table 1) since it led to the removal of as high as 88.4% of the cells in the `Kim2020` dataset. The valid numbers of cells per cell type in all four data sets are summarized (Supplementary Fig. 2). As part of pre-processing, all the locus-pairs along the diagonal of the contact matrices were excluded from the analysis. We observed distinct distributions of interaction frequencies among the diagonal and off-diagonal locus pairs (Supplementary Fig. 17) across the datasets. In the benchmarking experiments, locus-pairs at all genomic distances (excluding the diagonals) were utilized to avoid the exclusion of large percentages of interactions (Supplementary Fig. 18), except for scHiC Topics, which only focuses on locus-pairs within 10Mb.

We used existing bulk Hi-C datasets of specific cell types as the gold standard for comparing different methods. Specifically, GM12878 combined in situ and IMR90 combined in situ data were downloaded from GEO under accession GSE63525^35^. For differential interaction detection, two deeply sequenced biological replicates (replicates HIC019 and HIC020 for GM12878 and HIC051 and HIC056 for IMR90) were utilized. One combined replicate of HAP1 was obtained from GEO with the accession number GSE74072^41^ and another replicate utilized the wild-type condition data from GEO with accession number GSE95015^42^. H1ESC (accession 4DNESRJ8KV4Q) and HFF (accession 4DNES2R6PUEK) data were obtained from the 4D Nucleome portal^43^.

### Evaluation metrics

We considered both K-means clustering with and without low-dimensional projections with t-SNE and UMAP, and Louvain graph clustering^26^, with known numbers of clusters, to identify cell types. Application without t-SNE and UMAP involved applying K-means and Louvain graph clustering on the vectorized cell by locus-pairs matrices (for CellScale, BandScale, and BandNorm) or the method latent component embeddings. Applications with t-SNE and UMAP low-dimensional projections first leveraged PCA to reduce the dimensions of the vectorized cell by locus-pairs matrices or method latent component embeddings to the top 50 principal components before applying t-SNE and UMAP to generate low-dimensional embeddings. We quantified the resulting clustering performances with adjusted rand index (ARI)^44^ which measures the similarity between two data clusterings, i.e., true underlying cell types and the estimated clustering, adjusted for chance similarity. We also evaluated average silhouette scores^45^ to measure the separation between known cell types in the t-SNE and UMAP visualizations. Collectively, these led to six evaluation settings (Fig. 1C).

### Implementation details

#### scHiCluster

We customized the scHiCluster script from https://github.com/zhoujt1994/scHiCluster to accelerate the data loading procedure before running scHi-Cluster. The restart probability was set to 0.5 and the binarization percentile to 0.8. Following the scHiCluster application^15^, we chose the top 50 principal components for each chromosome, concatenated them, and applied another round of PCA to generate the top 50 principal components as the resulting low-dimensional embedding.

#### scHiC Topic

Following the pre-filtering guidelines from scHiC Topics^14^ (https://github.com/khj3017/schic-topic-model), only locus-pairs within 10Mb of each other were utilized for cell type separation. To determine the optimal topic number based on the silhouette score as suggested in the method paper^14^, we set the lowest topic number to be 10 and the highest to be 90 with increments of 5. The final optimal number of topics selected for `Ramani2017, Li2019` and `Kim2020` are 35, and 60 for `Lee2019`, respectively.

#### Higashi

All the parameters were set to defaults suggested by the Higashi pipeline (https://github.com/ma-compbio/Higashi) with the exception of `neighbor_num` which we set to 3, using the release version in January 2021 with commit ID 4f2ce9db4967f264042f052c38577f36d9f53681. We compared the performance of Higashi using GPU to only using CPU, therefore the `cpu_num` was set to 23 and `gpu_num` was set to 1 or 0 accordingly.

#### BandScale+CNN

The CNN was implemented with some modifications to the CNN Variational Autoencoder module of Pytorch (https://github.com/sksq96/pytorch-vae). Both the encoder and decoder are symmetric and contain two layers. The latent vector *z* was set to have 20 dimensions and `kernel_size` parameter to 4. Furthermore, `bias` was set to False and `stride` was set to 2 for all the layers. The minimal iteration parameter was set to 90 with a batch size of 10. The learning rate for the Adam algorithm was set to 1e-3.

#### 3DVI

The implementation of 3DVI on each band was based on the scVI-tools^17^ (https://github.com/YosefLab/scvi-tools), where we used the default parameters and set the latent variable dimension to 100. We filtered cells that have no interaction for all the locus-pairs in each band matrix. When concatenating the latent embeddings across band matrices, the latent value for the missing cells was filled with zero.

#### SVA

We first divided the obtained matrix by its minimum value to get a non-negative matrix and then use the *ComBat_seq* function in R package *sva* to do batch removal.

#### Seurat

We constructed the *ScaleData* and utilized *vars.to.regress* function in R package *Seurat* to regress out the batch effect.

#### Harmony

Use *HarmonyMatrix* function in R package *harmony* with *do_pca* = *FALSE* to remove the batch of the matrix.

#### diffHic

Differential chromatin interaction detection requires at least two replicate per condition. For differential detection between two aggregated scHi-C cell types, we randomly partition the cells with the same cell label into two groups, with each forming a pseudo-replicate. Differential detection is implemented using R package *diffHic*.

#### Evaluation metrics

K-means replied on the `kmeans` function of `stats` R package using the default `Hartigan-Wong` algorithm. `nstart` was set to 20 and `iter.max` to 1000. Louvain graph clustering was carried out based on `FindNeighbors` and `FindClusters` functions in `Seurat` R package. Silhouette coefficients were obtained using `silhouette` function from `cluster` R package.

#### De-noising performances with aggregated scHi-C data using cell labels from unsupervised clustering

We repeated the assessment of the impact of the normalization and de-noising methods on the downstream analysis using the cell type labels inferred from clustering in aggregating the scHi-C contact matrices. This ensured a more unbiased assessment of the overall effect of the analysis methods without relying on true cell type labels (Supplementary Fig. 19). K-means clustering of the UMAP embeddings of the `Kim2020` dataset resulted in four clusters which we labeled as GM12878, H1ESC, HAP1, and HFF cell type (Supplementary Fig. 19A). Concordant with the results that relied on true cell labels, detection of A/B compartments and topologically associating domains (TADs), contact matrix similarity, detection of interacting locus-pairs yield the advantages of BandNorm followed by 3DVI (Supplementary Fig. 19B-E). Similarly, comparison with respect to the differentially interacting locus-pairs resulted in similar performances across the methods (Supplementary Fig. 19F-G).

## Code availability

R package, BandNorm, together with the curated scHi-C datasets are available at https://github.com/keleslab/BandNorm. 3DVI pipeline is implemented in Python in a way that allows parallelization in high-performance computing environments with source codes and instructions available at https://github.com/keleslab/3DVI.

