## Supplementary Table and Figures for "Normalization and De-noising of Single-cell Hi-C Data with BandNorm and 3DVI"

Supplementary Table 1: Summary of single-cell Hi-C data.

| Data Source | Cell Type | # of Cell | Interaction Frequency | Off-diagonal Interaction Frequency | # of locus-pairs | # of Off-diag locus-pairs | Mean Off-diagonal Interaction Frequency per Cell | Mean # of Off-diagonal locus-pairs per Cell |
|---|---|---|---|---|---|---|---|---|
| Ramani2017 | GM12878, HAP1, Hela, K562 | 2,610 | 51,420,688 | 23,089,064 (44.9%) | 16,639,155 | 11,081,243 (66.6%) | 8,846.38 | 4,245.69 |
| Lee2019 | Astro, Endo, L23, L4, L5, L6, MG, MP, Ndnf, ODC, OPC, Pvalb, Sst, Vip | 4,238 | 4,586,887,008 | 553,896,738 (12.08%) | 132,310,725 | 120,126,414 (90.79%) | 130,821.2 | 28,371.85 |
| Li2019 | 2i, Serum1, Serum2 | 150 | 15,846,298 | 2,612,886 (16.49%) | 1,906,405 | 1,523,307 (79.9%) | 17,419.24 | 10,155.38 |
| Kim2020 | GM12878, H1Esc, HAP1, HFF, IMR90 | 9,230 | 95,764,991 | 39,702,592 (41.46%) | 48,866,268 | 30,333,591 (62.07%) | 4,305.2 | 3,289.26 |

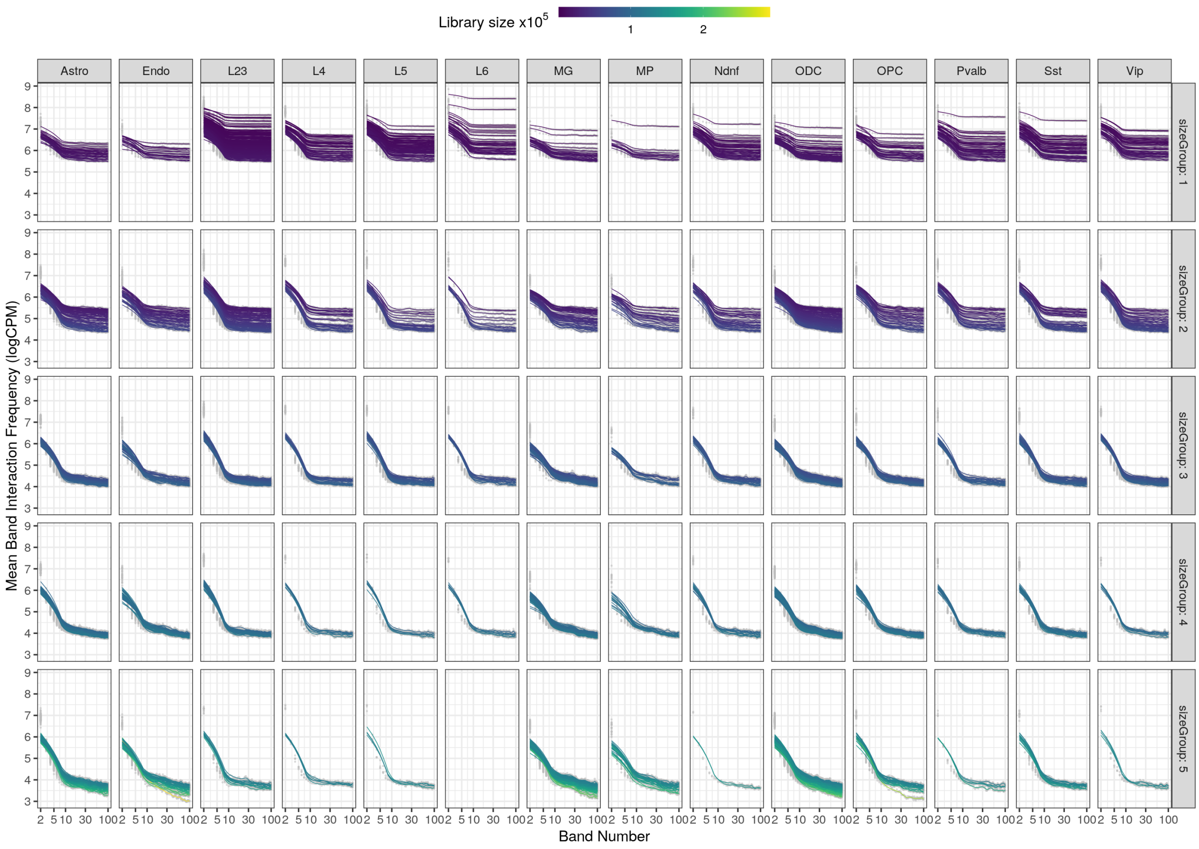


**Supplementary Figure 1** *Genomic distance effect at different library size levels.* Band effect on the mean band interaction frequencies as a function of library size and cell type for the `Lee2019` dataset with 14 cell types. Each line depicts the lowess smoothed mean of the interaction frequencies (in log counts per million) with respect to the genomic distance, i.e., the band number, from individual cells.

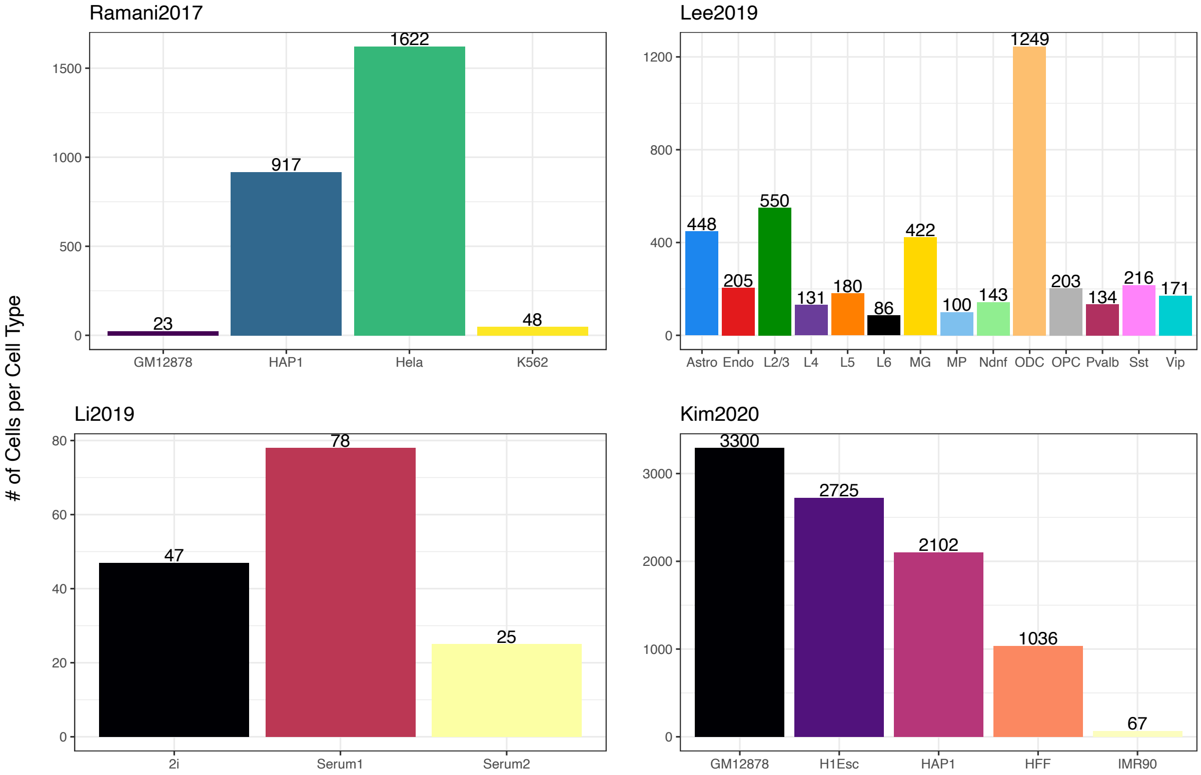


**Supplementary Figure 2** *Number of valid cells for each cell type in the* `Ramani2017`, `Lee2019`, `Li2019`, and `Kim2020` *datasets.* Details of cell filtering are available in the Methods.

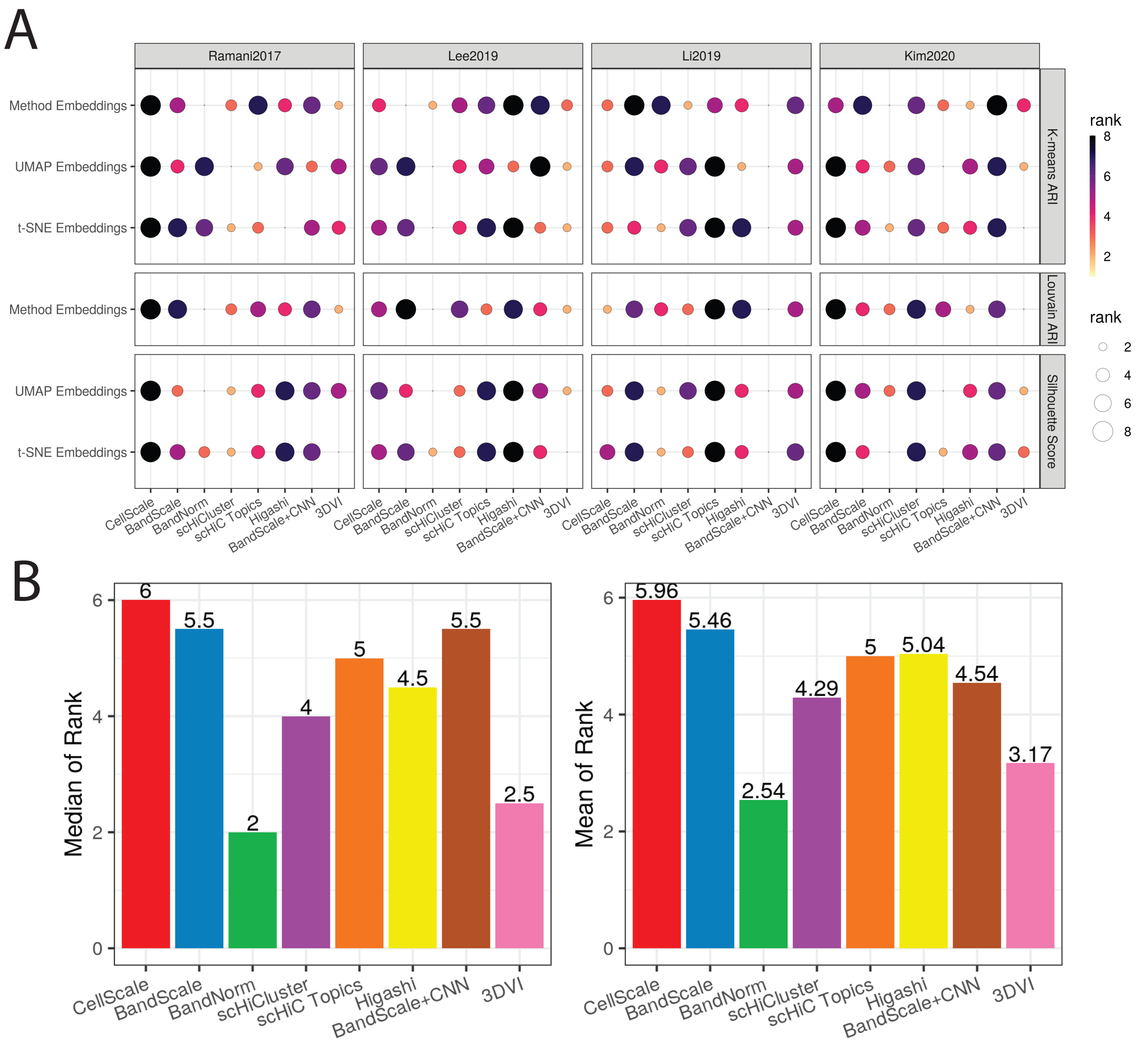


**Supplementary Figure 3** *Performance ranking of the scHi-C normalization and denoising methods.* **A.** Rank of the eight scHi-C analysis methods, CellScale, BandScale, BandNorm, scHiCluster, scHiC Topics, Higashi, BandScale + CNN, and 3DVI, across the six evaluation metrics and four benchmark data sets. **B.** Median and mean ranks of the scHi-C methods across the six evaluation metrics and four data sets.

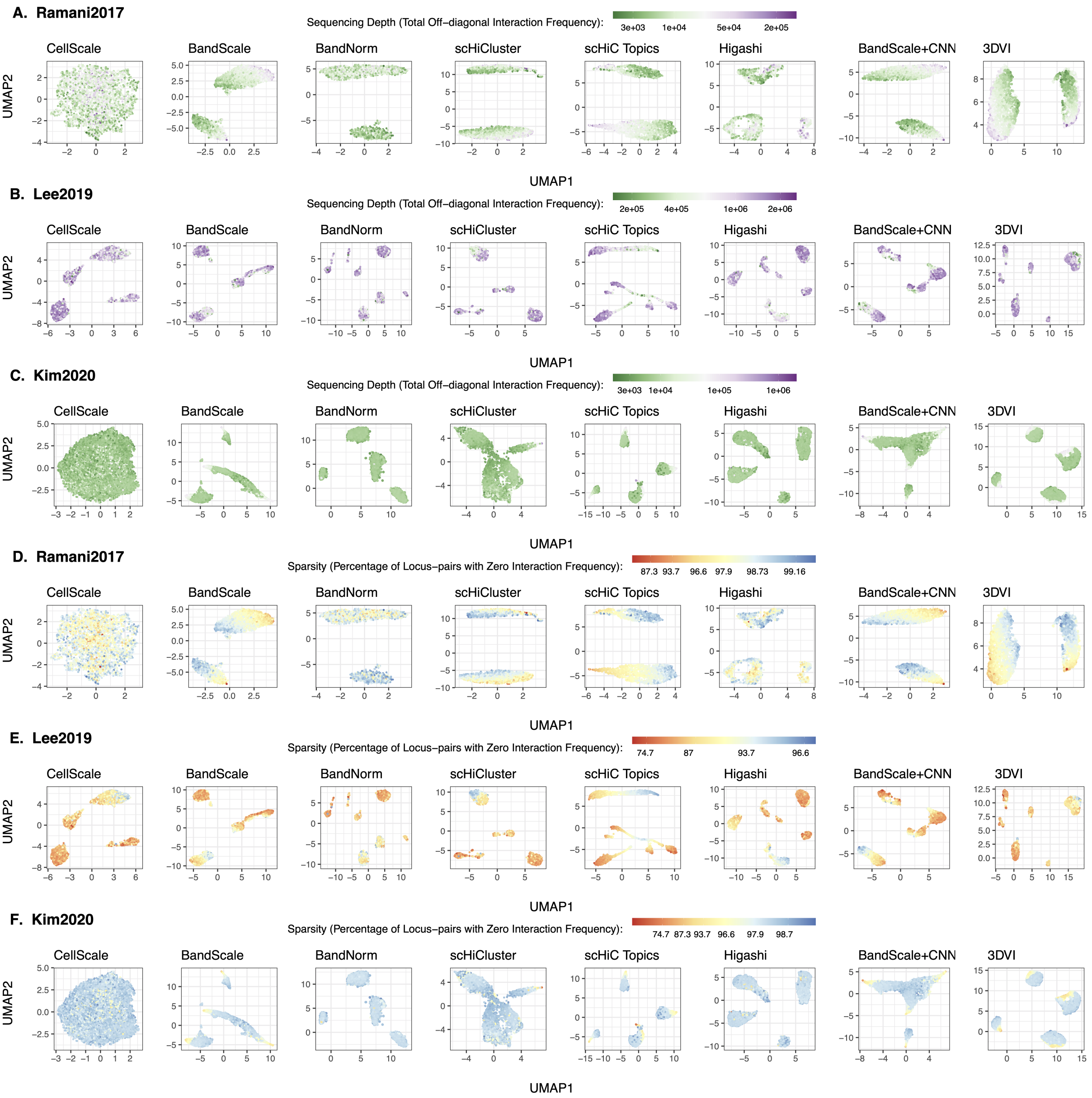


**Supplementary Figure 4** *Impact of sequencing depth and sparsity on cell type separation.* Sequencing depth for **A.** `Ramani2017`, **B.** `Lee2019` and **C.** `Kim2020` data sets are computed as the total interaction frequencies in the upper triangular of the contact matrices. The results are displayed using scatter plots of the two UMAP coordinates. Color shading depicts the sequencing depths of the cells. Sparsity for **D.** `Ramani2017`, **E.** `Lee2019` and **F.** `Kim2020` data sets is defined as the percentage of locus-pairs with zero interaction frequency in the upper triangular of the contact matrix. The results are displayed using scatter plots of the two UMAP coordinates. Color shading depicts the sparsity of the cells.

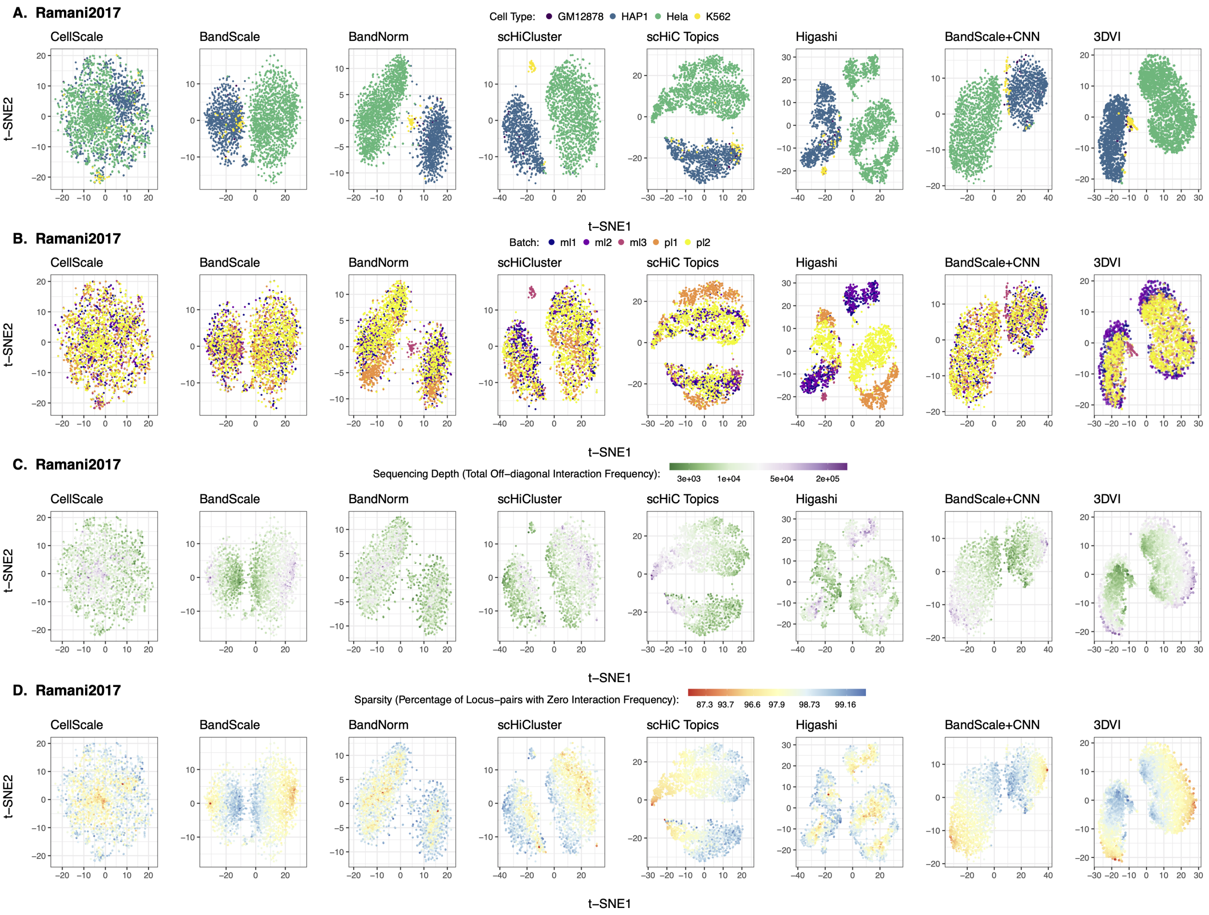


**Supplementary Figure 5** *t-SNE visualization of the low-dimensional embeddings of the `Ramani2017` data set.* Cell type separation performance (**A**), and the impact of batch effect (**B**), sequencing depth (**C**) and sparsity (**D**) for eight scHi-C normalization and de-noising methods. The results are displayed using scatter plots of the two t-SNE coordinates.

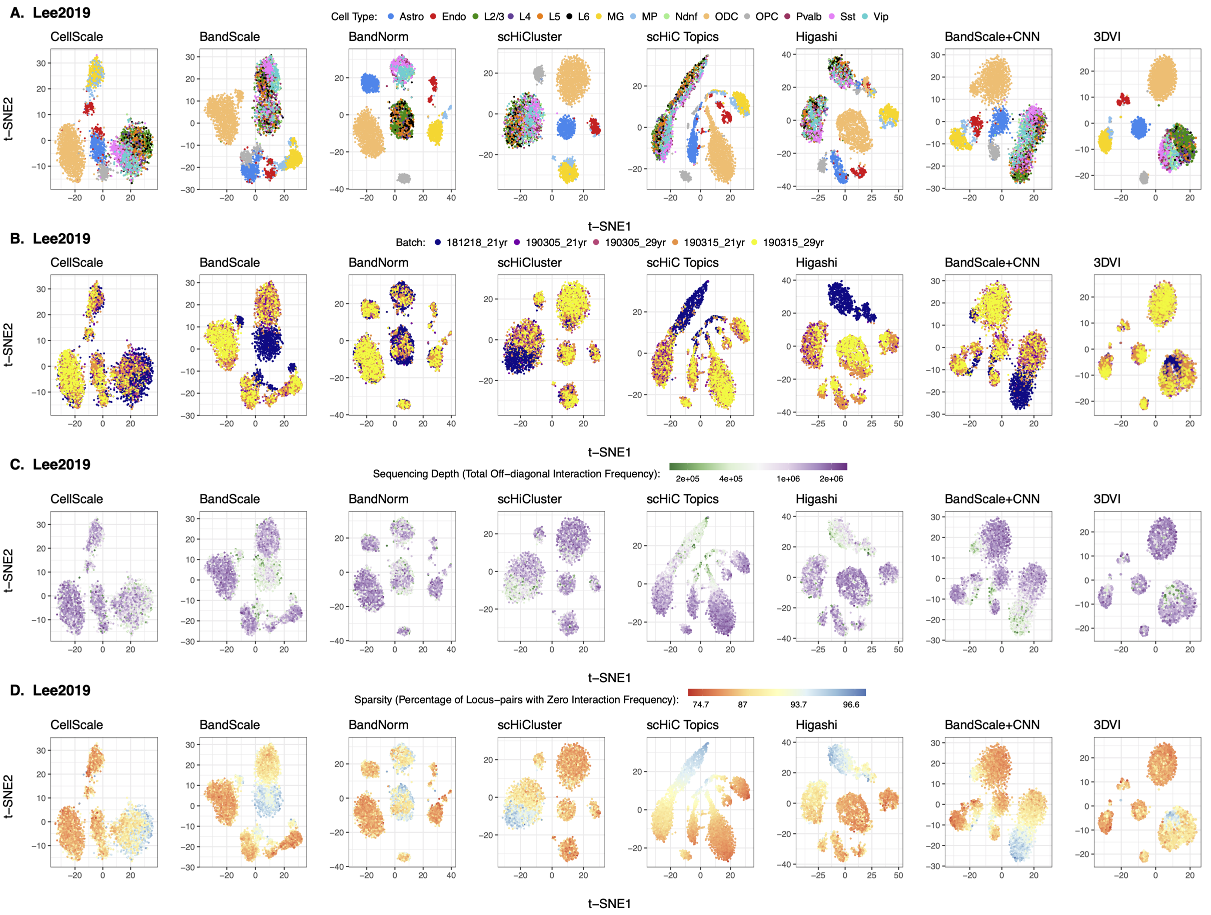


**Supplementary Figure 6** *t-SNE visualization of the low-dimensional embeddings of the `Lee2019` data set.* Cell type separation performance (**A**), and the impact of batch effect (**B**), sequencing depth (**C**) and sparsity (**D**) for the scHi-C normalization and de-noising methods. The results are displayed using scatter plots of the two t-SNE coordinates.

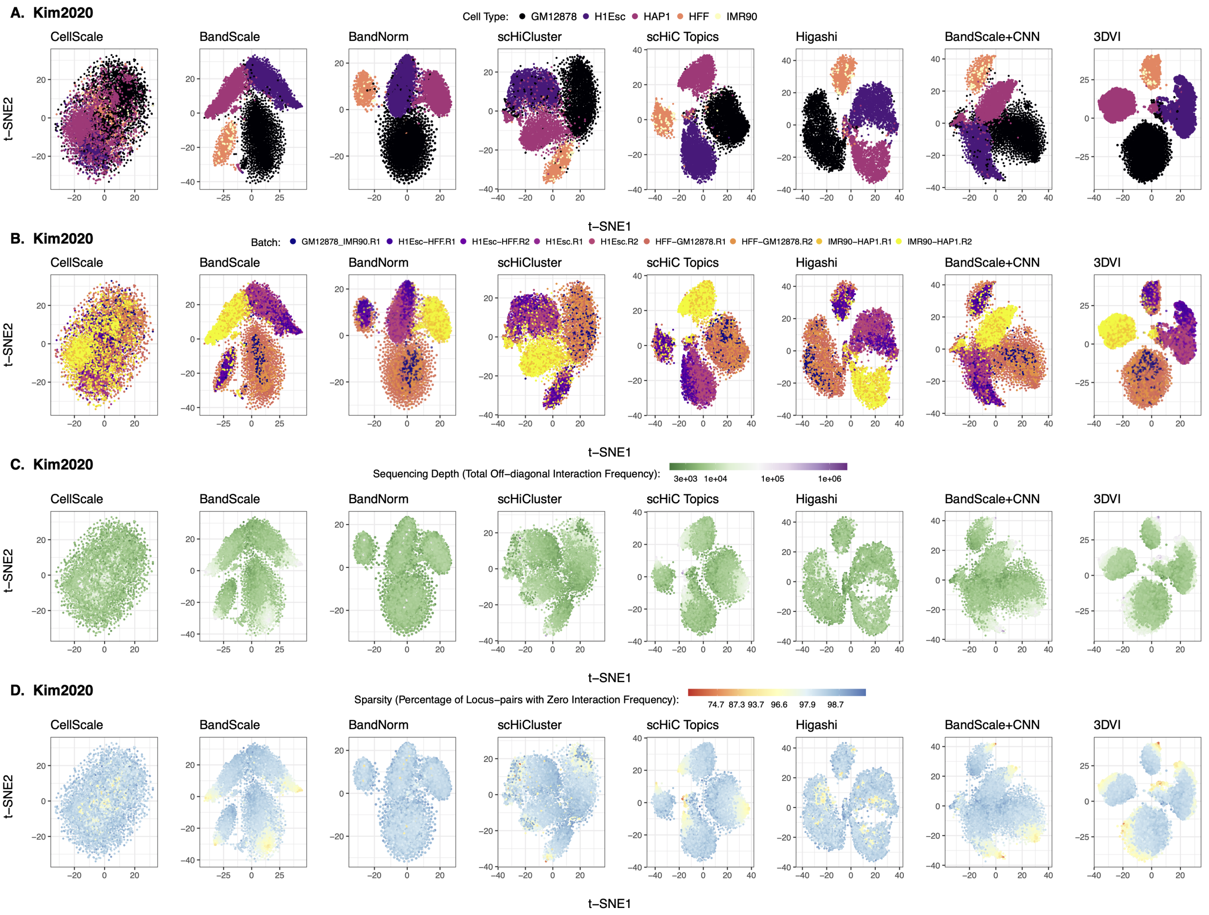


**Supplementary Figure 7** *t-SNE visualization of the low-dimensional embeddings of the* `Kim2020` *data set.* Cell type separation performance (**A**), and the impact of batch effect (**B**), sequencing depth (**C**) and sparsity (**D**) for the scHi-C normalization and de-noising methods. The results are displayed using scatter plots of the two t-SNE coordinates.

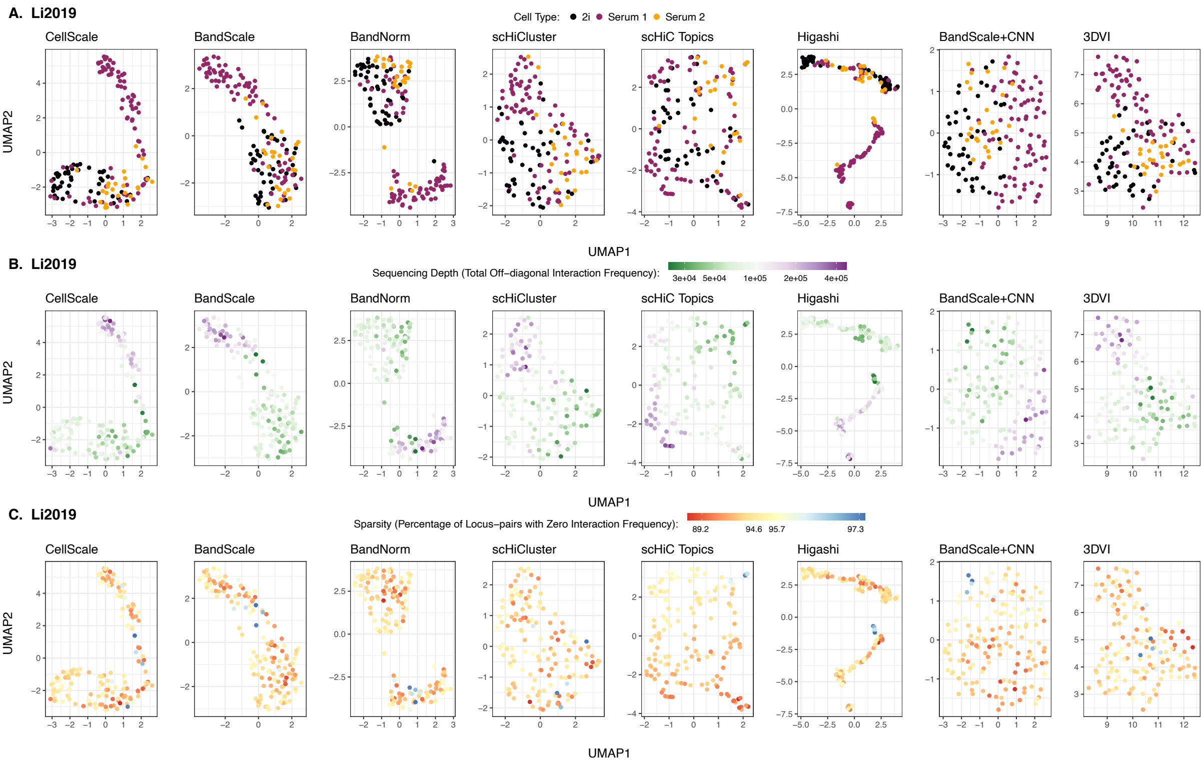


**Supplementary Figure 8** *UMAP visualization of the low-dimensional embeddings of the* `Li2019` *data set.* Cell type separation performance (**A**), sequencing depth (**B**) and sparsity (**C**) for the scHi-C normalization and de-noising methods. The results are displayed using scatter plots of the two UMAP coordinates.

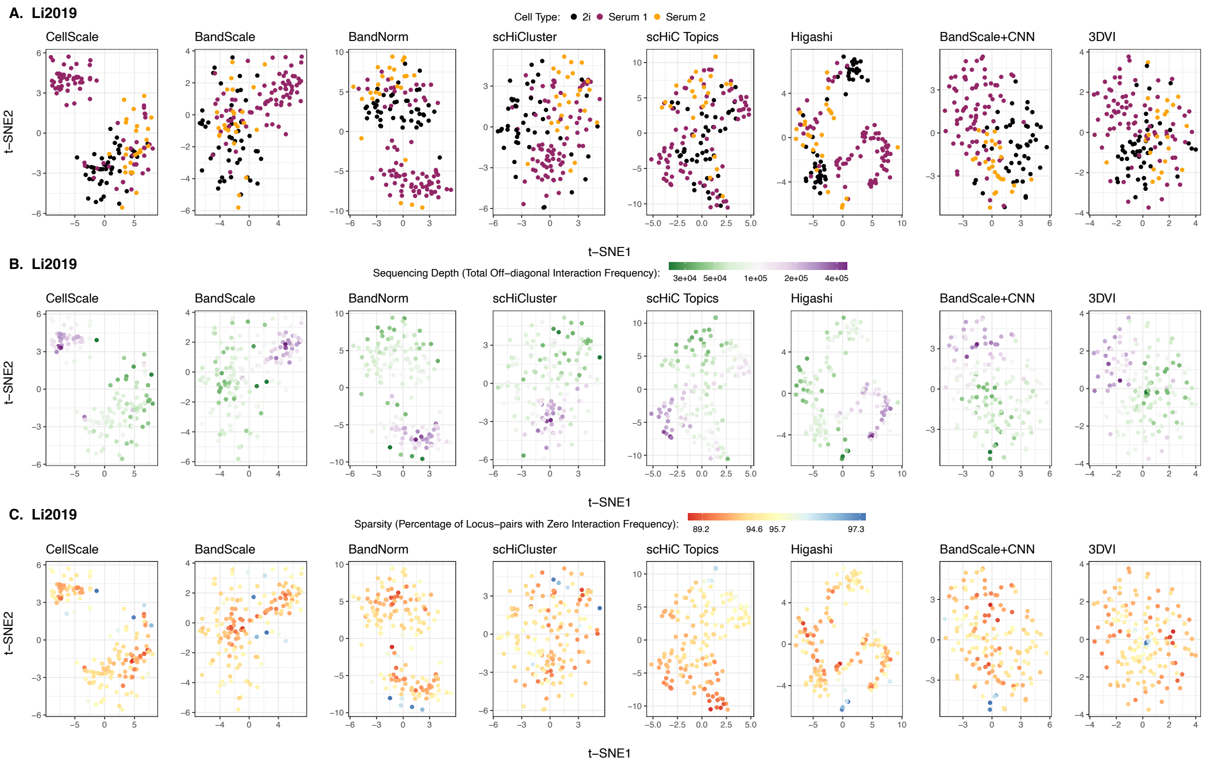


**Supplementary Figure 9** *t-SNE visualization of the low-dimensional embeddings of the `Li2019` data set.* Cell type separation performance (**A**), sequencing depth (**B**) and sparsity (**C**) for the scHi-C normalization and de-noising methods. The results are displayed using scatter plots of the two t-SNE coordinates.

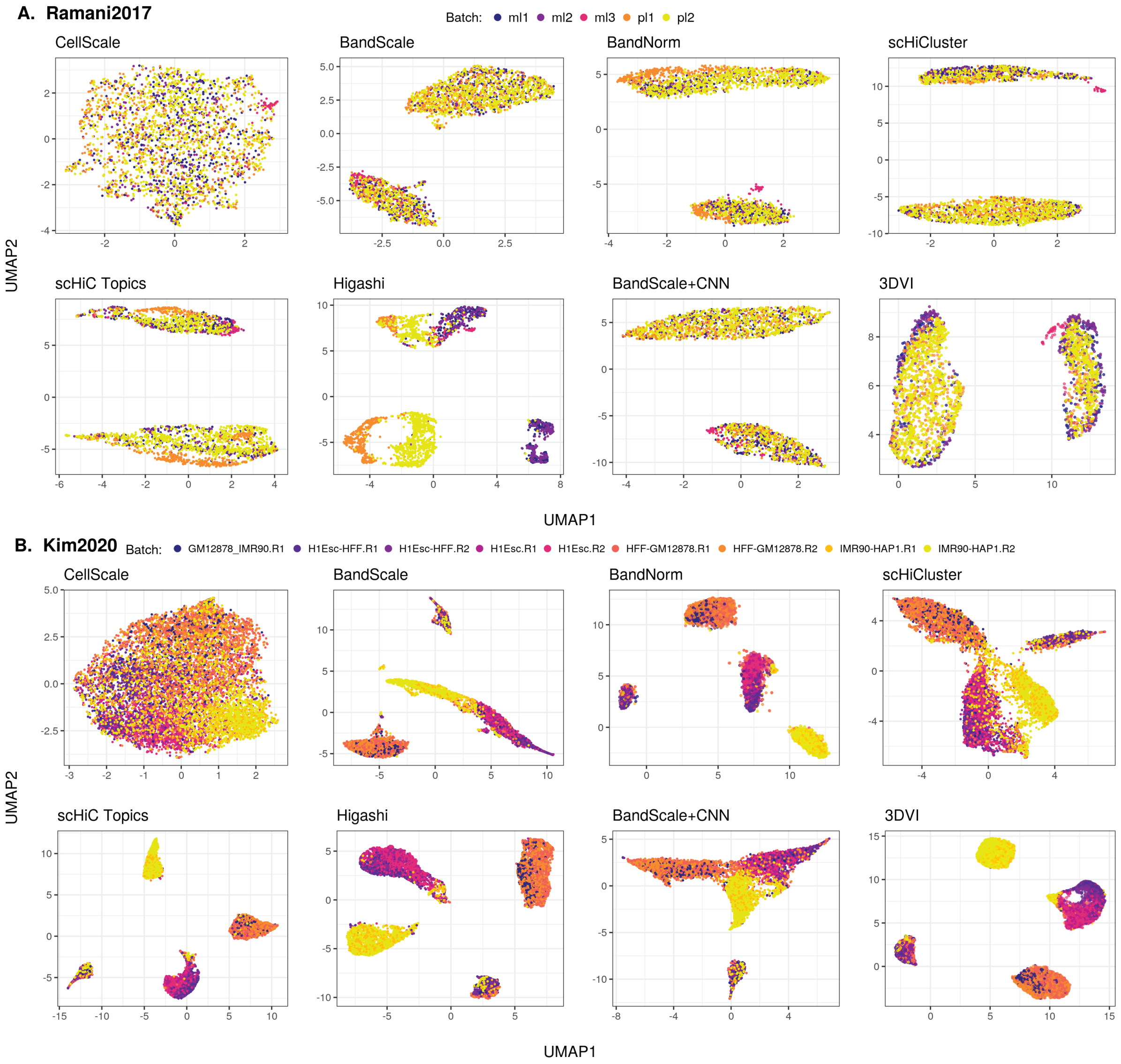


**Supplementary Figure 10** *Batch effect on cell type separation.* Batch effect on the cell type separation for the scHi-C normalization and de-noising methods using `Ramani2017` (**A**) and `Kim2020` (**B**) data sets. The results are displayed using scatter plots of the two UMAP coordinates. Color shading depicts the batches of the cells.

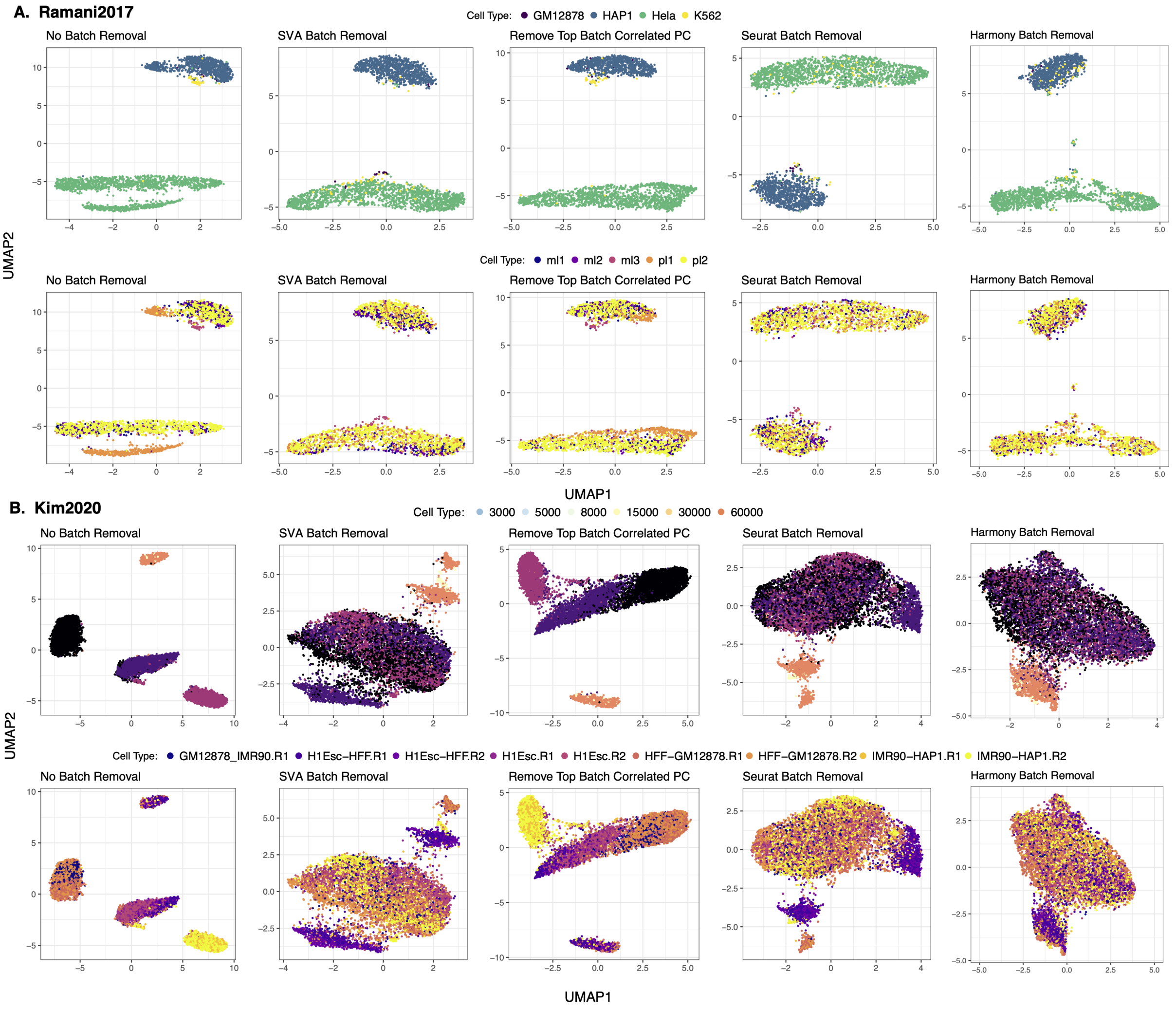


**Supplementary Figure 11** *Comparison of batch effect removal methods on scHi-C data after BandNorm normalization.* Performance of four batch effect removal methods, SVA[27], removing the top correlated principal component, Seurat batch effect regression[28], and Harmony[29], applied with BandNorm normalization on `Ramani2017` and `Kim2020` data sets.

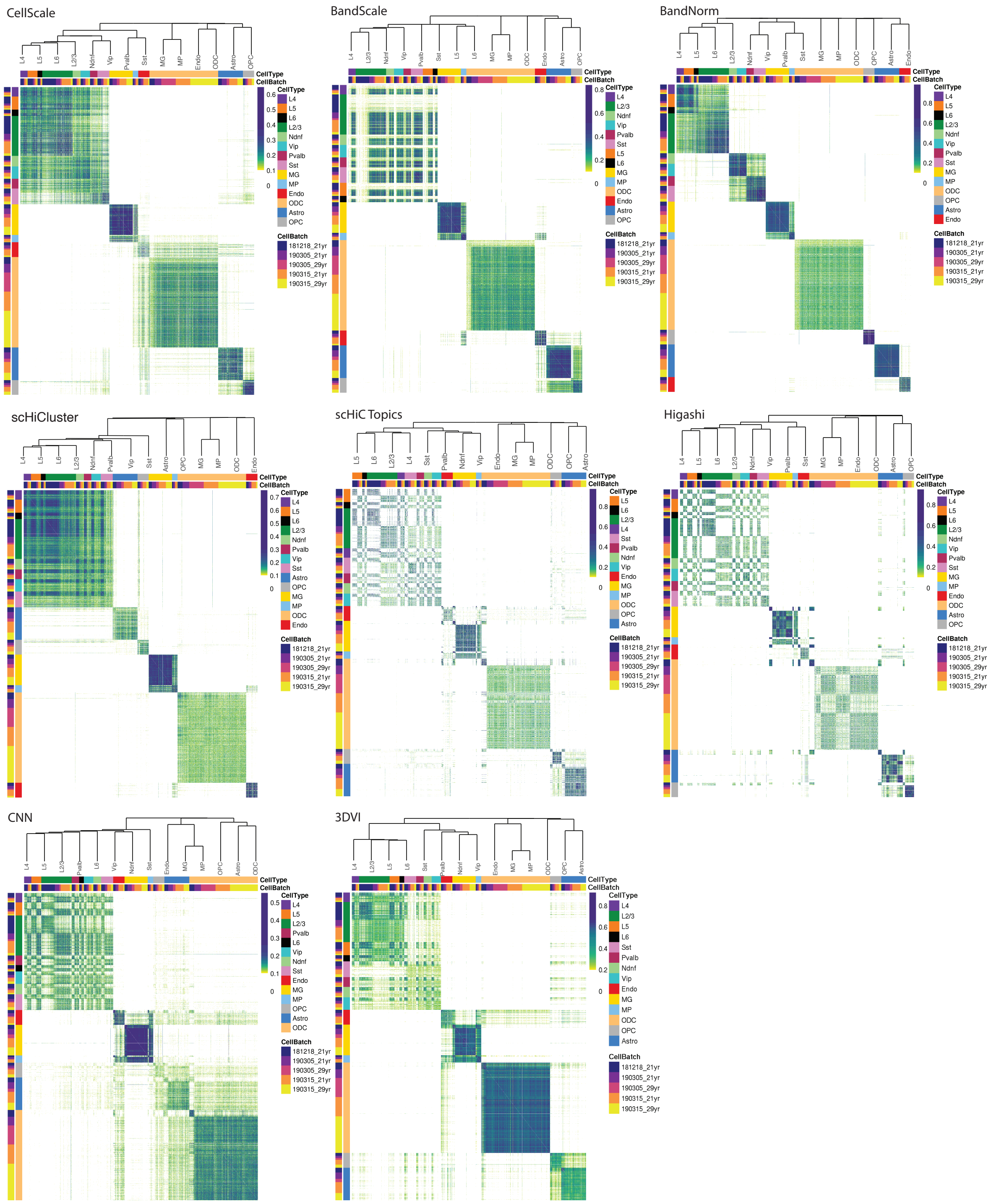


**Supplementary Figure 12** *Pairwise cell similarities based on low-dimensional embeddings to elucidate cell type relationships.* Pairwise similarities between the cells are computed based on the low-dimensional embeddings of the scHi-C data by each method. The similarity scores are the edge weights constructed based on the shared nearest neighbors graphs of `Lee2019` data set to elucidate the cell type relationships and demonstrate batch effects.

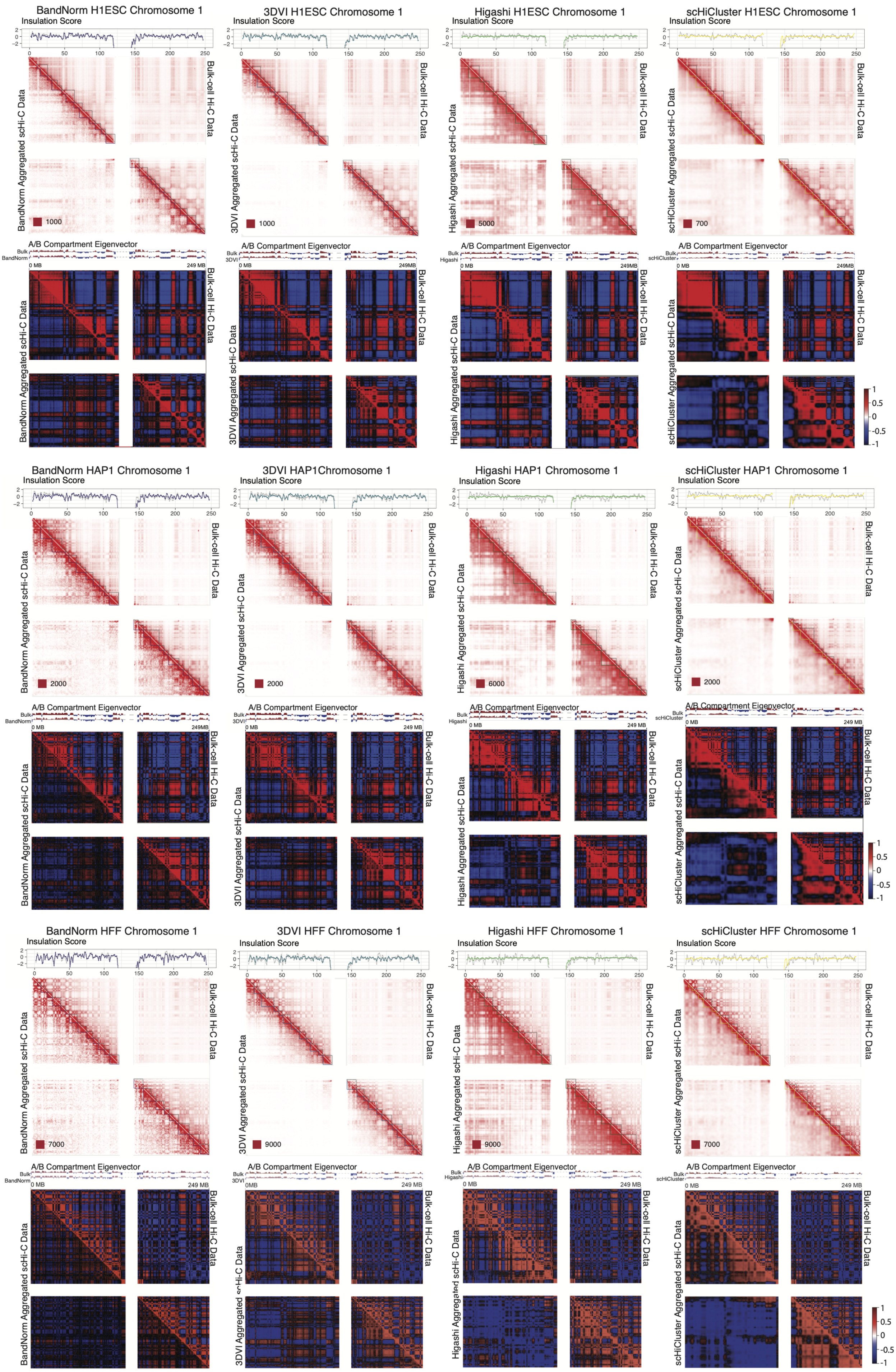
BandNorm H1ESC Chromosome 1
3DVI H1ESC Chromosome 1
Higashi H1ESC Chromosome 1
scHiCluster H1ESC Chromosome 1
Insulation Score
A/B Compartment Eigenvector
Bulk-cell Hi-C Data
BandNorm Aggregated scHi-C Data
3DVI Aggregated scHi-C Data
Higashi Aggregated scHi-C Data
scHiCluster Aggregated scHi-C Data
BandNorm HAP1 Chromosome 1
3DVI HAP1Chromosome 1
Higashi HAP1 Chromosome 1
scHiCluster HAP1 Chromosome 1
BandNorm HFF Chromosome 1
3DVI HFF Chromosome 1
Higashi HFF Chromosome 1
scHiCluster HFF Chromosome 1

**Supplementary Figure 13** *Evaluation for the detection of topologically associating domains (TADs) and A/B compartments in H1ESC, HAP1, and HFF cells.* Comparison of detected TADs and A/B compartment between bulk Hi-C data (upper right triangles) and the aggregated single-cell Hi-C data (lower left triangles) after normalization or de-noising on `Kim2020` data set with known H1ESC, HAP1, and HFF cell type labels. The insulation scores[46] that trace the TAD boundaries are depicted above the contact matrices with grey lines corresponding to bulk Hi-C data and purple for BandNorm, blue for 3DVI, green for Higashi, and yellow for scHiCluster. The numbers after the red squares at the left bottom of each contact matrix represent the minimum interaction frequency for the reddest locus-pair. A/B compartments are detected using the eigenvector of correlation map of bulk (upper right triangles) or aggregated (lower left triangles) Hi-C matrices, values of which are displayed above each correlation matrix.

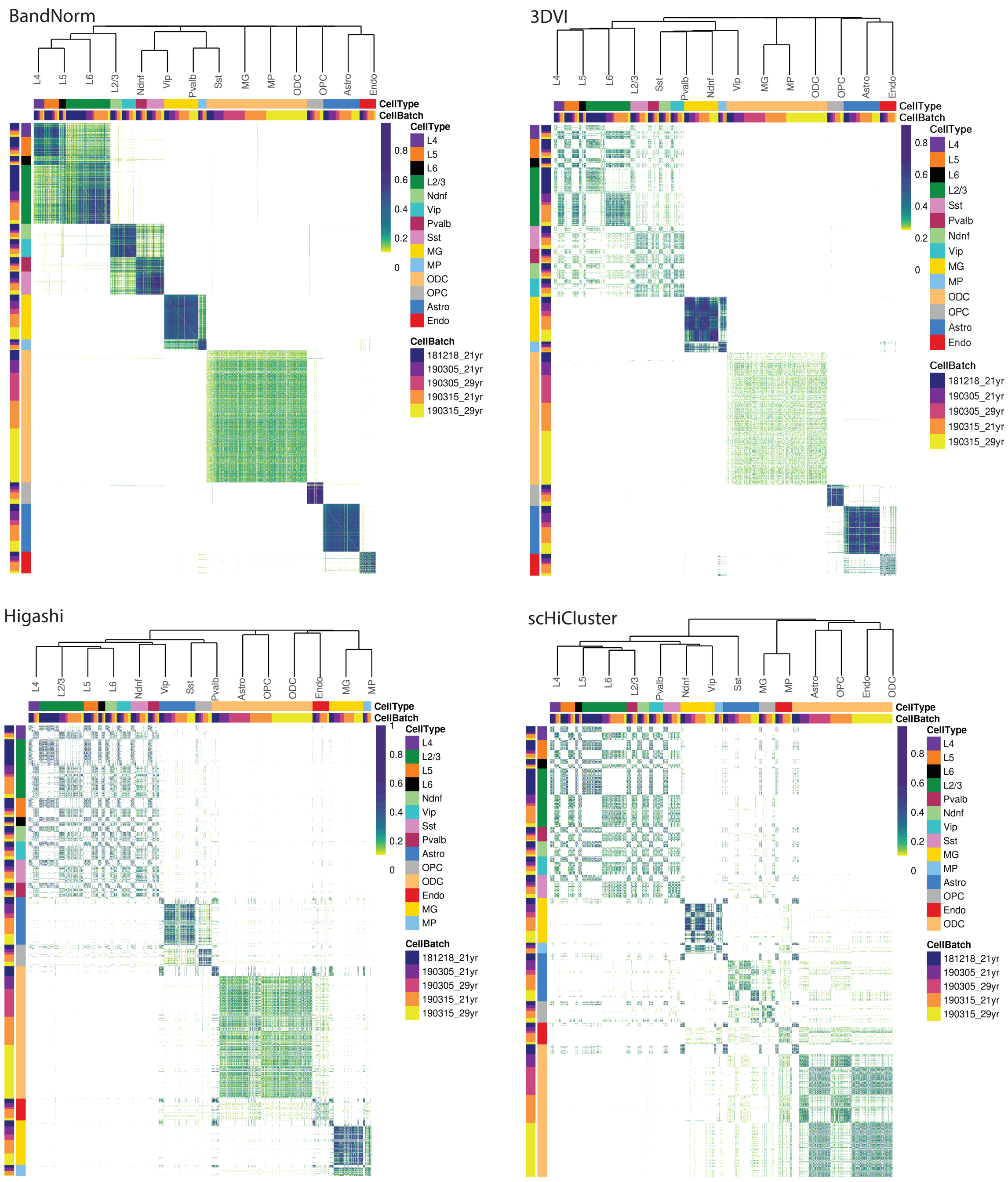


**Supplementary Figure 14** *Similarity scores between aggregated scHi-C matrices and the corresponding bulk Hi-C matrices.* The scores are constructed with the edge weights of shared nearest neighbors graph of `Lee2019` data set to depict the cell type relationships and demonstrate batch effect.

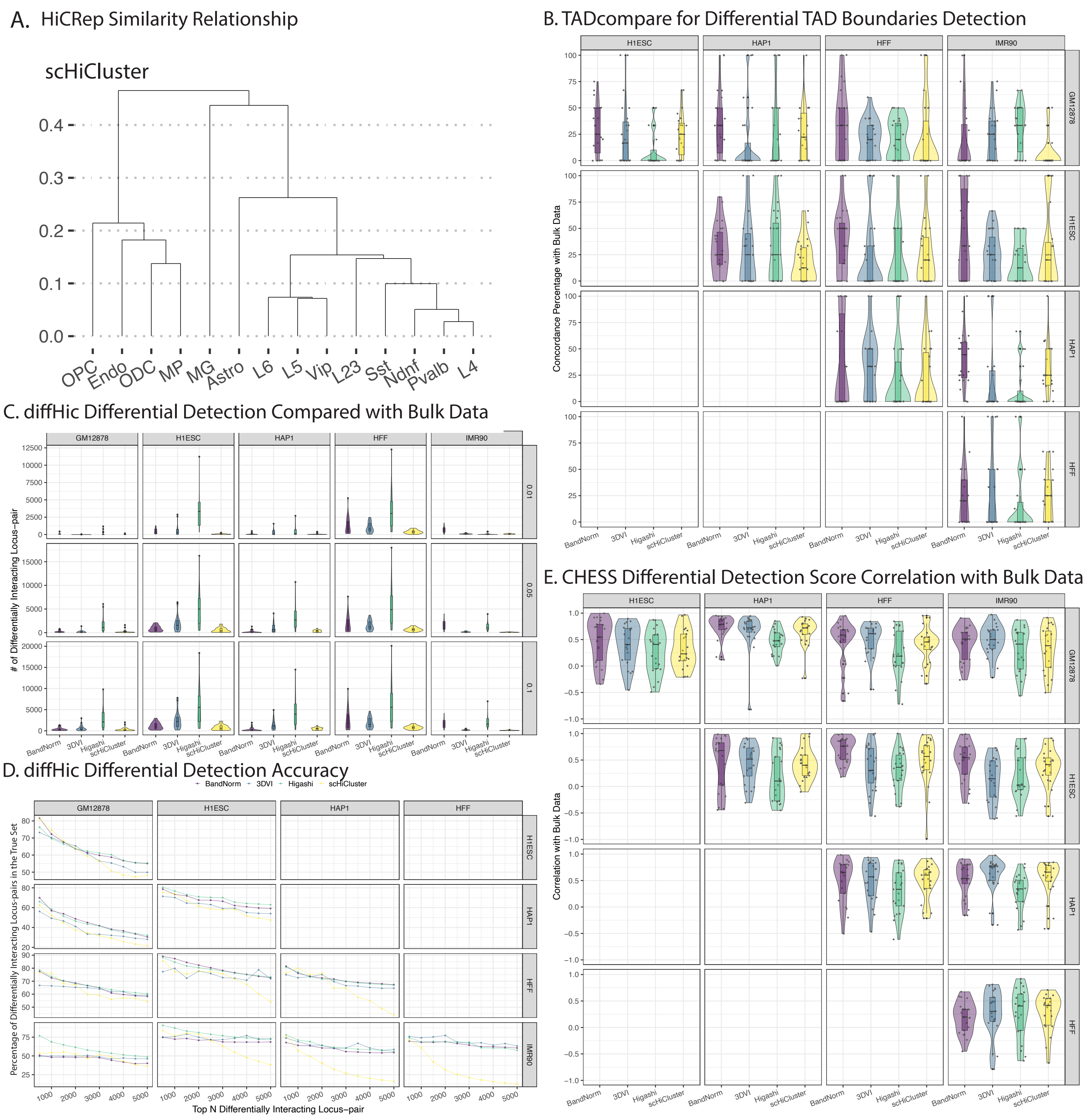


**Supplementary Figure 15** *Domain and locus-pair differential analysis of aggregated scHi-C data after normalization and de-noising, with known true cell type labels.* **A.** Hierarchical clustering depicting cell type relationship is constructed using the HiCRep[12] similarity scores between every pair of the aggregated scHi-C cell type after de-noising with scHiCluster. **B.** Comparison of differential TAD boundaries detected by

TADcompare[32] between every pair of cell types. **C.** Comparison of differentially interacting locus-pairs detected between bulk and aggregated scHi-C after normalization or de-noising, using diffHic[33]. The three rows correspond to filtering of the differential chromatin interactions at three FDR levels, i.e., adjusted $P$ value $\leq 0.01$ for the first row. **D.** Percentage of top N (N = 500, 1,000, ...) significant differentially interacting locus-pairs, detected by diffHic[33] analysis of aggregated scHi-C matrices from each method, that are in the gold standard set. The gold standard set is defined as the significant differentially interacting locus-pairs detected by diffHic[33] from the cell type specific bulk Hi-C data. **E.** Correlation of CHESS[34] scores, depicting differential interactions of the cell types, between bulk Hi-C and aggregated scHi-C from different methods. Sample sizes for each violin plot of **B, C, E** are n = 23 corresponding to 23 chromosomes investigated.

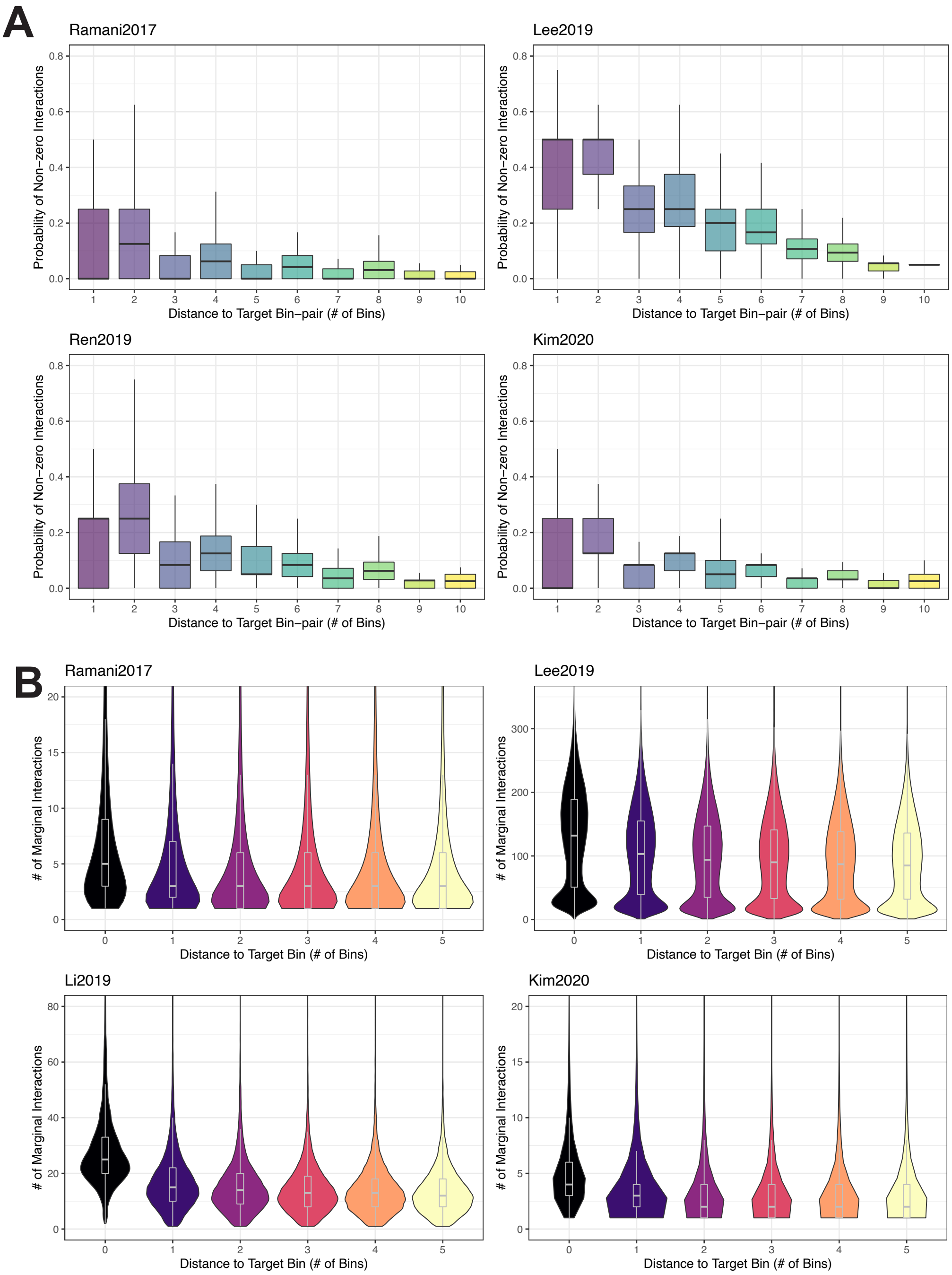


**Supplementary Figure 16** *Neighborhood effect at 1Mb resolution.* **A.** Empirical probability of non-zero interaction around the neighboring locus-pairs of top 10 locus-pairs with interaction frequencies (referred to as high IF locus-pairs) $\geq 3$ on each chromosome contact matrix per cell. Distance is defined as the number of locus-pairs between the neighboring locus-pair and the high IF locus-pair. **B.** The genomic loci are first ordered based on their marginal interactions (row sum of the contact matrices) and the top 10 loci for each cell are considered per chromosome per cell. Boxplots depict the interaction frequencies around these loci as a function of genomic distance measured in numbers of locus-pairs between the high IF locus-pairs and their neighbors.

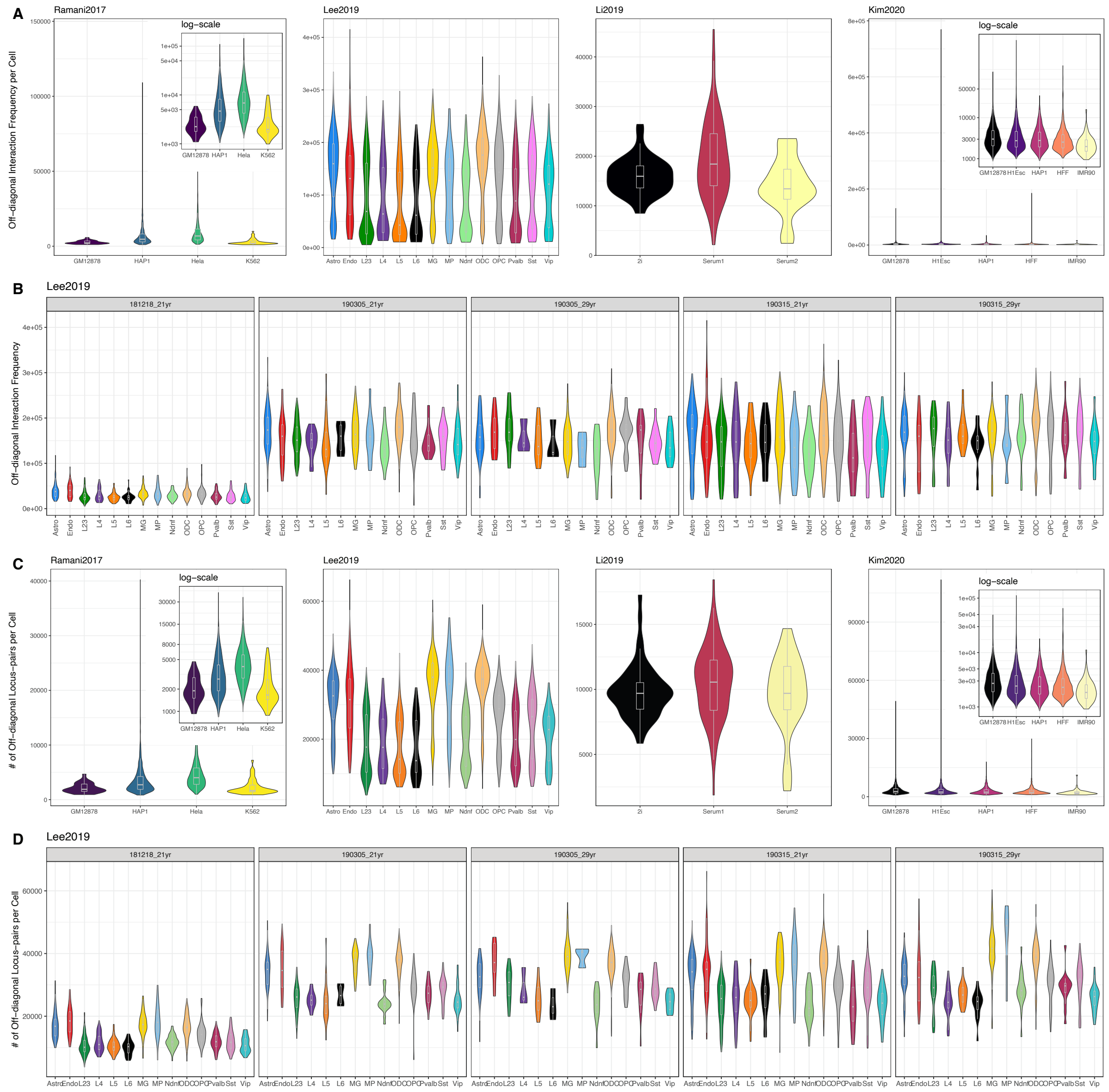


**Supplementary Figure 17** *Distribution of the total off-diagonal interaction frequencies and the numbers of non-zero off-diagonal locus-pairs per cell.* **A.** Distribution of the total interaction frequencies across the contact matrices (excluding the main diagonal) of the cells for each cell type of `Ramani2017`, `Lee2019`, `Li2019`, and `Kim2020` data sets. The small panels within the figure display the same data with the y-axis is in log-scale. **B.** Total interaction frequencies per cell for the `Lee2019` data set stratified by the five libraries. The cells from the 181218_21 yr batch have relatively lower interaction frequency leading to a bimodal pattern in the cell-level total off-diagonal interaction frequencies in **A**. **C.** Summary of the total numbers of locus-pairs with non-zero interaction frequencies (excluding the locus-pairs on the main diagonal) per cell for each cell type of `Ramani2017`, `Lee2019`, `Li2019`, and `Kim2020` data sets. The small panels within the figure display the same data with the y-axis is in log-scale. **D.** Total number of non-zero off-diagonal locus-pairs per cell for the `Lee2019` data set stratified by the five libraries. The sample sizes for each cell type violin plot of **A-D** per data set correspond to the valid cell number shown in Supplementary Fig. 2.

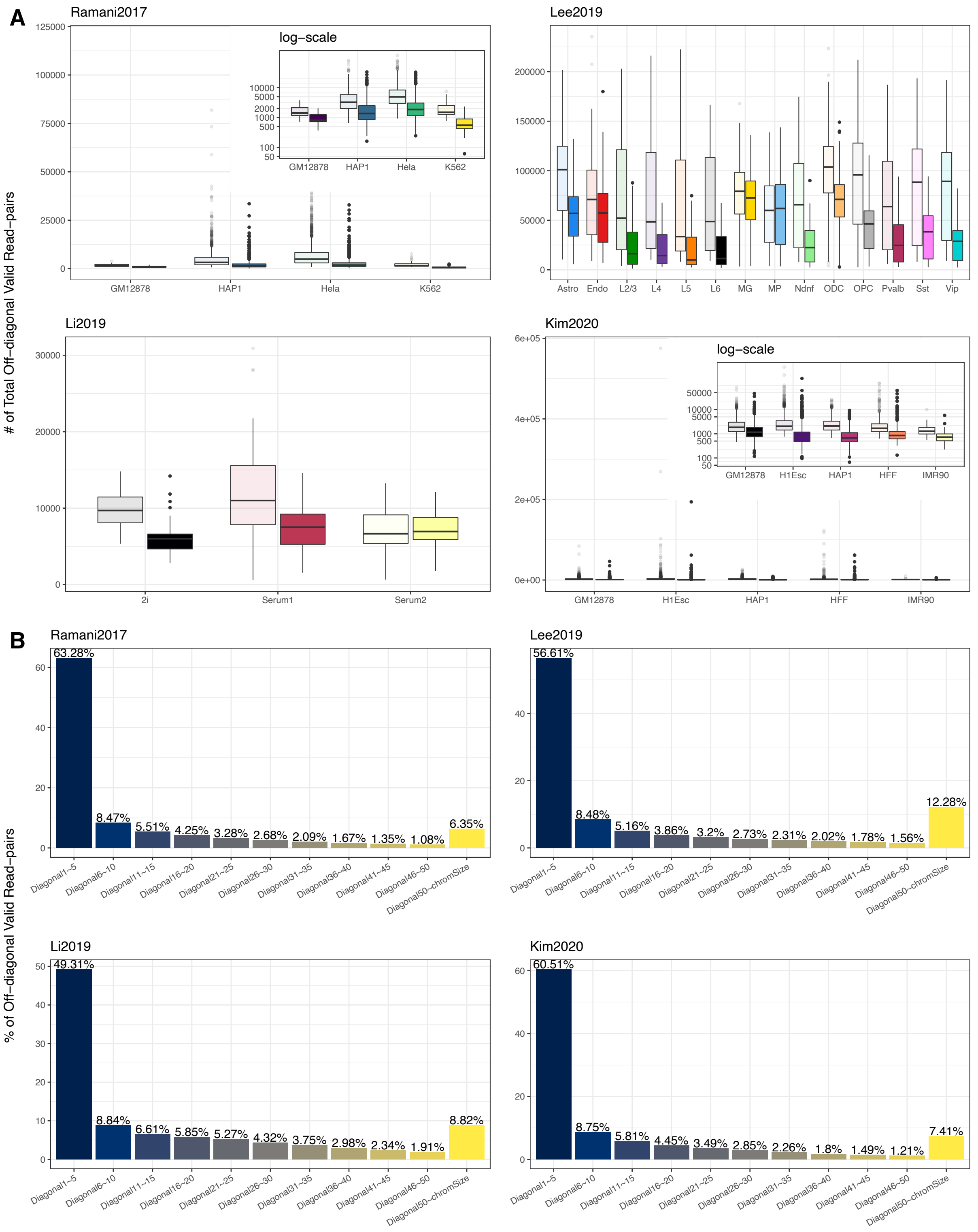


**Supplementary Figure 18** *Genomic distance impact on the interaction frequency.* **A.** Total off-diagonal interaction frequency per cell for each cell type stratified by the interaction distance. First boxplot for each cell type depicts interactions between locus-pairs that are within 10Mb of each other and the second box plot per cell type depicts interactions between locus-pairs that are separated by more than 10Mb. **B.** Percentage of total interactions at different band intervals, excluding the matrix diagonal.

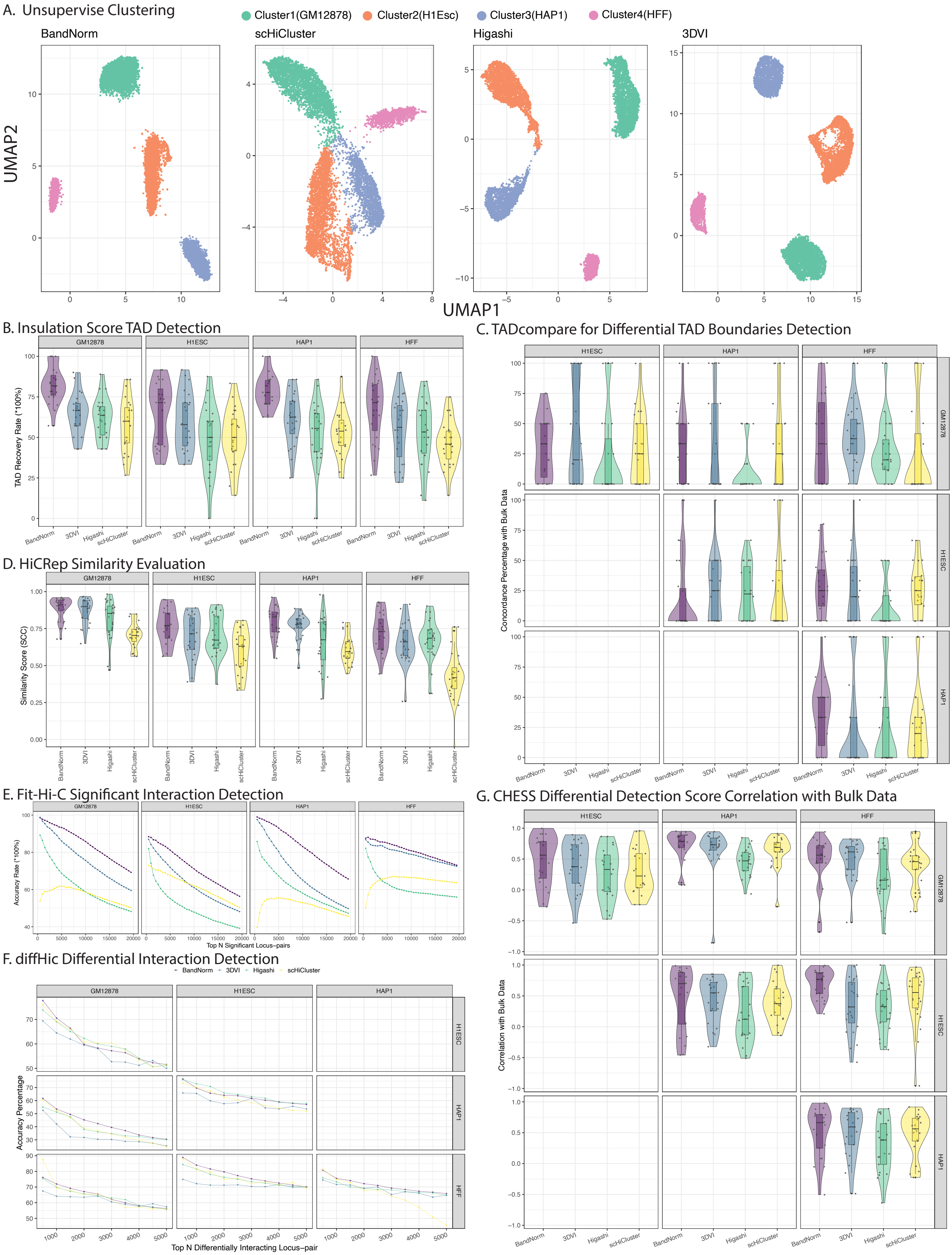
A. Unsupervise Clustering
Cluster1(GM12878)
Cluster2(H1Esc)
Cluster3(HAP1)
Cluster4(HFF)
BandNorm
scHiCluster
Higashi
3DVI
UMAP2
UMAP1
B. Insulation Score TAD Detection
TAD Recovery Rate (*100%)
C. TADcompare for Differential TAD Boundaries Detection
Concordance Percentage with Bulk Data
D. HiCRep Similarity Evaluation
Similarity Score (SCC)
E. Fit-Hi-C Significant Interaction Detection
Accuracy Rate (*100%)
Top N Significant Locus-pairs
F. diffHic Differential Interaction Detection
Accuracy Percentage
Top N Differentially Interacting Locus-pair
G. CHESS Differential Detection Score Correlation with Bulk Data
Correlation with Bulk Data
GM12878
H1ESC
HAP1
HFF

**Supplementary Figure 19** *Evaluation of impact on downstream analysis of Hi-C data with cell type labels inferred from unsupervised clustering.* **A.** K-means clustering on the UMAP projections of low-dimensional latent embeddings of BandNorm, scHiCluster, Higashi, and 3DVI. The clusters identified are labeled with cell types GM12878, H1ESC, HAP1, and HFF based on comparison with bulk Hi-C data. **B.** Percentage of TAD boundaries (based on Insulation Score[36]) that is within 1Mb distance of the corresponding bulk cell type Hi-C data TAD boundaries. **C.** Differential TAD boundaries detected by TADcompare[32] between every pair of clusters. **D.** HiCRep similarity of the aggregated scHi-C data of individual clusters with the bulk Hi-C data. **E.** Percentage of top N (N = 5,000, 10,000, ...) significant interacting locus-pairs that are in the gold standard set for each method. The gold standard set is defined as the top 50,000 significant locus-pairs detected by Fit-Hi-C[31] from the cell type specific bulk Hi-C data. **F.** Percentage of top N (N = 500, 1,000, ...) significant differentially interacting locus-pairs, detected by diffHic[33] analysis of aggregated scHi-C matrices from each method, that are in the gold standard set. The gold standard set is defined as the significant differentially interacting locus-pairs detected by diffHic[33] from the cell type specific bulk Hi-C data. **G.** Correlation of CHESS[34] scores, depicting differential interactions of the cell types, between bulk Hi-C and aggregated scHi-C from different methods. Sample sizes for each violin plot of **B-D, G** are n = 23 corresponding to 23 chromosomes investigated.
